# Transformer models of mutation risk at base-pair resolution identify non-coding hotspot cancer driver mutations

**DOI:** 10.64898/2026.07.07.736824

**Authors:** Iván Galván-Femenía, Marcell Veiner, Daniel Naro, Fran Supek

**Author notes:** Universitat Politècnica de Catalunya, Barcelona, Spain (Daniel Naro).

## Abstract

Recurrent somatic mutations reveal cancer drivers, but in whole genomes many non-coding hotspots are passengers generated by localized mutational processes. We developed MutFormer, a transformer/convolutional neural net model that predicts base-pair-resolution somatic mutation risk from DNA sequence alone, separately for COSMIC signatures. Trained on >90 million high-confidence mutational signature-assigned SNVs from cancer genomes, MutFormer learns extended sequence determinants beyond trinucleotide context, often spanning up to ~20 nucleotides, and recovers APOBEC, UV, POLE and SBS17 sequence preferences as well as various additional mutation risk-prone motifs. We integrated MutFormer predictions with mutation burden, signature exposures and epigenetic covariates to model neutral recurrence of individual hotspots in >18,000 tumor whole genomes. Coding-region analyses calibrated the framework against known driver genes and AlphaMissense scores, supporting conservative false-discovery estimates. In non-coding regions, most recurrent hotspots were explained by passenger mutability, whereas selected outliers were enriched near cancer genes and supported by SpliceAI, PromoterAI, AlphaGenome and expression data. Prioritized candidates include splice-region or deep-intronic hotspots in *BCL6*, *PTEN*, *TCF7L2*, *PBRM1*, *PTPRT* and *VHL*, and promoter hotspots in *SHKBP1*, *PRSS3* and *BCL2*.

## Introduction

Cancer genomes are shaped by the combined action of mutational processes and natural selection^1–3^. Most somatic mutations are passengers, accumulated as a consequence of DNA damage, replication errors and imperfect repair^4,5^. A small minority are drivers that alter cellular fitness and are therefore recurrently selected during tumor evolution; distinguishing these two classes is a central problem in cancer genomics^6,7^. Recurrence is a powerful signal of selection, and it underlies the discovery of many canonical driver mutations in coding regions. However, recurrence alone is not sufficient: a mutation may recur because it is beneficial to the tumor, or simply because the affected nucleotide is unusually mutable^7–10^.

This ambiguity is especially acute in the non-coding genome^11,12^. Whole-genome sequencing has enabled systematic searches for driver mutations in promoters, untranslated regions, enhancers, splice-regulatory elements and non-coding RNA genes. Yet, despite various efforts, robustly supported non-coding drivers remain rare, with *TERT* promoter mutations standing as the clearest example^1^. Several studies have shown that many apparently compelling non-coding hotspots are instead explained by localized mutational processes, technical artifacts, or extreme context-dependent mutability. UV-associated mutations accumulate at regulatory motifs in skin cancers^8,13,14^, APOBEC enzymes generate recurrent mutations in susceptible DNA structures^10,15,16^, CTCF/cohesin-bound sites can show elevated mutability in gastrointestinal and skin tumors^17,18^, and somatic hypermutation affects specific regulatory loci in lymphoid malignancies^19^. Thus, the central challenge is not merely to find recurrent non-coding mutations, but to determine whether their recurrence exceeds what should be expected from the local mutational baseline.

Somatic mutation rates vary across the genome at many spatial scales^20^. At the megabase scale, mutation density is associated with replication timing, chromatin state and nuclear organization^21–24^. At gene scale, transcription modulates local mutation rates via transcription-coupled repair, or in certain cases by transcription-associated mutagenesis^25–28^. At subgenic scale, regulatory factor occupancy, chromatin accessibility or local hypomethylation can generate focal hyper- or hypomutation^13,29–32^. At the finest scale, the immediate nucleotide neighborhood is a major determinant of mutability. Mutational signatures summarize this process-specific variation in the form of characteristic substitution spectra, usually defined by the mutated base and its two flanking nucleotides^33–35^. This trinucleotide representation has been highly successful for identifying the mutagenic processes active in tumors, but it is also a compression of the sequence information available around each mutation. Many mutational processes in human somatic or germline cells are known, or suspected, to depend on sequence features extending beyond the trinucleotide context^3,36–40^.

Recent work has begun to reveal this hidden layer of local mutability. Extended k-mer analyses have shown that hotspots often mark highly mutable sequence contexts, and that signatures such as UV, POLE proofreading deficiency, APOBEC activity and SBS17-like processes have sequence dependencies broader than the conventional trinucleotide^8,15,17,41^. Other studies have built genome-wide mutation-rate models using epigenetic features, local mutation density, nucleotide content or predefined genomic annotations, enabling driver searches in arbitrary regions or sliding windows^2,4,5,42,43^. These approaches have been instrumental in demonstrating the importance of rigorous background models. However, non-coding driver analyses still test short regions rather than individual bases, model mutation rates typically at kilobase or larger resolution, and treat sequence context using fixed k-mer features. As a result, they are not optimized to answer a critical question: what is the expected recurrence of this exact nucleotide substitution, in this cancer type, given the mutational processes active in the tumor and the extended local sequence in which the mutation occurs?

Deep learning provides an opportunity to address this gap. Neural net sequence models have transformed regulatory genomics by learning complex motif grammar directly from DNA^44,44–46^, predicting gene activity and chromatin state, and related approaches have been used to model fine-scale germline mutation rates^47–49^. In somatic cancer genomes, however, a base-pair-resolution model must account for the fact that different mutational processes obey different sequence rules. A local DNA sequence that is highly mutable under UV damage may not be highly mutable under APOBEC activity, mismatch repair deficiency, POLE proofreading deficiency or SBS5 clock-like mutagenesis. Therefore, a useful model of local passenger mutation risk should be process-specific, nucleotide-resolved, and capable of learning complex sequence features.

Here, we introduce MutFormer, a transformer-based neural network model that predicts local somatic mutation risk from DNA sequence alone. MutFormer is trained separately for individual COSMIC SBS signatures, using matched unmutated control sites with the same trinucleotide context as each mutated site. This design forces the model to learn sequence determinants beyond the conventional trinucleotide representation of mutational signatures. Across 41 SBS signatures extracted from 12,166 whole cancer genomes, MutFormer reveals that many mutational processes depend on extended local sequence context, often spanning approximately 20 nucleotides. Model interpretation by *in silico* mutagenesis and motif discovery recovers known APOBEC, UV, POLE and SBS17-associated sequence preferences, while also showing that extended sequence context-dependent mutability is a widespread property of somatic mutagenesis by various signatures.

We then use MutFormer predictions as a process-aware mutational baseline for hotspot discovery in 18,317 whole cancer genomes (Supp Fig 1). By estimating the expected recurrence of individual nucleotide hotspots under neutrality, our framework distinguishes selected hotspots from passengers generated by extreme local mutability. Coding-region analyses validate the approach against known driver genes and AlphaMissense variant-impact scores. Applying the same single-base-resolution framework to promoters, UTRs, enhancers, splice regions and non-coding RNA genes shows that most recurrent non-coding hotspots are compatible with passenger mutability, but identifies a restricted set of candidate regulatory and splice-altering drivers supported by orthogonal functional predictions.

## Results

### Mutational signature extraction and assignment in a large cancer WGS collection

In our studies of local determinants of mutation risk, we separated mutations by the trinucleotide mutational signature, so we could model the risk incurred by the DNA extended context separately for each mutagenic process. Mutational signatures were extracted, using the non-negative matrix factorization (NMF) algorithm implemented in SigProfiler tool^50^, for 12,166 WGS of tumors containing ~230M somatic single-nucleotide variants (SNVs) and ~45M somatic indels (IDs). The matrices of SBS96 and ID83 counts were concatenated together as input for NMF, as this approach^51^ showed higher stability of the signatures than using the SBS96 matrix alone (Supp Fig 2). We considered 46 signatures an appropriate solution for a pancancer analysis (average silhouette index = 0.81), broadly agreeing with contemporaneous large-scale studies, which confirmed 42 of the COSMIC v3.4 catalog signatures^33^ in additional data and analyses^52^. Of our 46 signatures, we further focussed on the SNV spectrum, and used SigProfiler to fit them to the 41 known SBS mutational signatures in the COSMIC v3.4 catalog. Then, for each SNV mutation, we assigned it to the SBS mutational signature that is most likely to have caused that mutation, accounting for the mutation’s trinucleotide context and the activity of the SBS signatures in the genome containing the mutation (similar as described^53^, see Methods); finally we discarded the mutations where the assignment to signature was ambiguous. A total of ~93M SNVs with high-confidence assignments were classified into 41 different COSMIC SBS signatures, and further analyzed separately to assess local DNA sequence determinants of mutation rate at basepair resolution, for each mutational signature.

### MutFormer: a transformer neural network model of mutation rate at base-pair scale

There exist known examples where the extended DNA nucleotide context has effects on mutation risk, such as APOBEC-hypermutable sites with sequences forming DNA secondary structures^15^, the SBS17 and UV hypermutable sites at loci containing CTCF binding motifs^14,54^, UV-caused hypermutation at sites with ETS binding motifs at promoters^14,55^. Generalizing over these examples, we asked if neural networks (NNs) could be employed to search for DNA sequences, outside of the trinucleotide, that modulate local mutation risk of various mutagenic mechanisms. Motivated by the success of the transformer architecture in NN that predict gene regulation from DNA sequence^46,56^, wherein the self-attention mechanism of the transformer is thought to facilitate modeling interactions between various positions in DNA, we also implemented a combination of transformers with convolutional NNs for mutation risk modeling. Our NN inputs DNA sequences of 50 nucleotides upstream and downstream from the mutated site (Fig 1a). It uses a series of convolutional blocks with skip connections which enables the network to learn local DNA motifs, followed by multi-headed attention to learn relevant motif interactions (see Methods). This block is followed by a final sequential head and a sigmoid layer that compares the set of mutated sequences (encoded as class 1) with a set of non-mutated sites with matching trinucleotide context (encoded as class 0) that were sampled from the 1kb neighboring region of the mutated sites (Fig 1b). We optimized the NN architecture to achieve robust generalizability, i.e. ability to predict mutation risk on held-out data, across diverse mutational signatures. To this end, we employed a Bayesian hyperparameter optimization^57^ implemented in Weights & Biases to identify a NN configuration, performing a 25-iteration search on datasets consisting of SBS1, SBS2, SBS10a, and SBS17b signatures and the matched unmutated sites (Supp Fig 4). For each of the 4 selected signatures, the two top-performing parameter sets were then trained on the remaining 3 signatures, resulting in a performance ranking for each NN configuration (Supp Fig 5). Finally, we selected configuration #6, hereafter referred to as MutFormer, due its consistent out-of-sample performance across the 4 mutational signatures and relatively light-weight complexity. MutFormer contains a total of 5,901,825 trainable parameters, distributed in 3 repeated convolutional blocks (with kernel sizes 7, 3, and 3), and one multi-head attention block (8 heads) with rotary positional encoding and using flash attention. Throughout the blocks, the number of channels is kept at a constant 512 (Methods).

**Figure 1.**
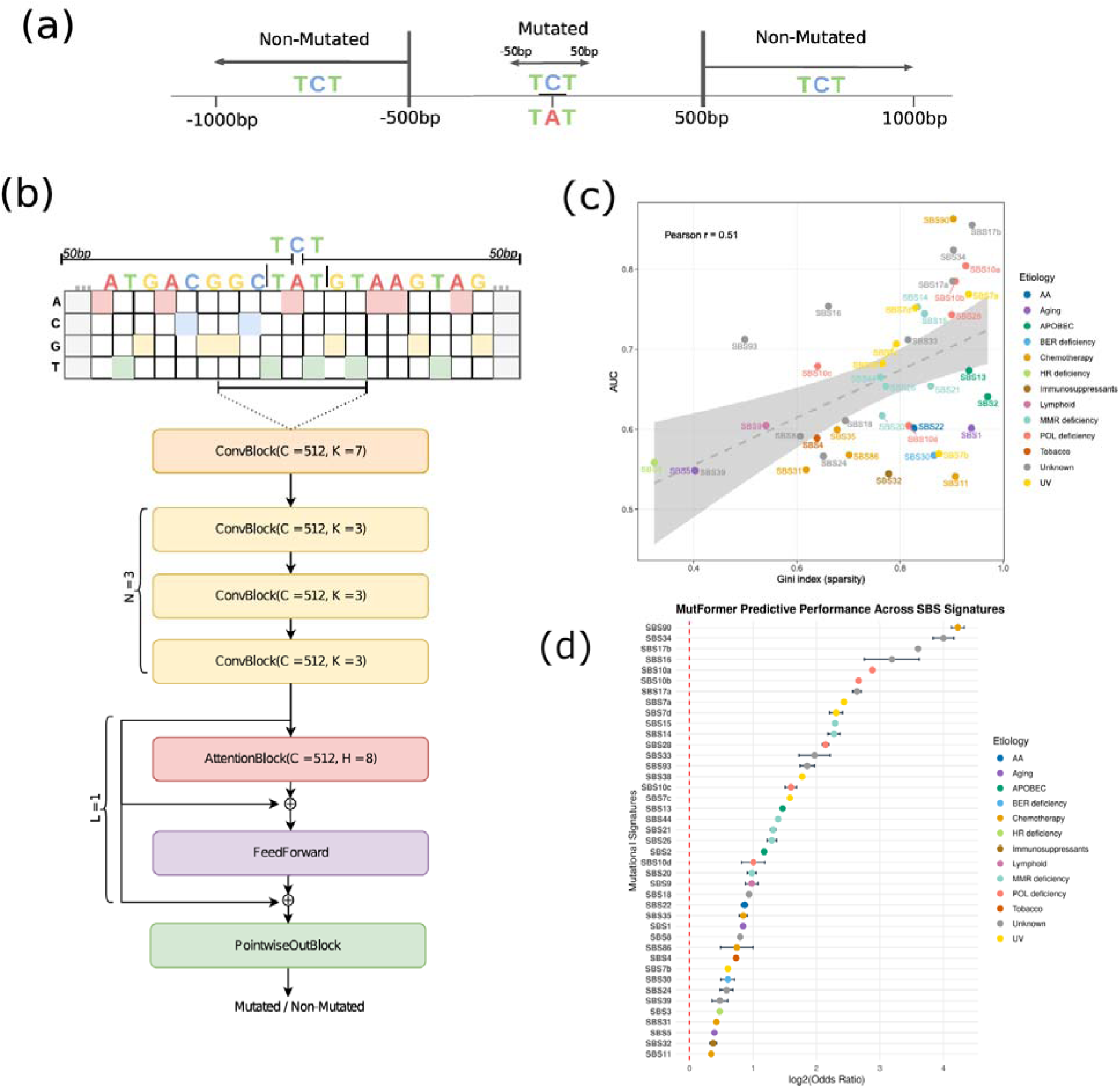
MutFormer neural network architecture and training scheme for modeling nucleotide-resolution, extended context-dependent mutation risk. **(a)** Mutated sites affected by a particular SBS signature are matched with non-mutated control sites with the same trinucleotide context, sampled from the surrounding ±1 kb genomic region excluding a 500 bp buffer region. MutFormer discriminates mutated sites based on extended nucleotide context, up to +-50bp, beyond the trinucleotide. **(b)** Overview of the convolutional–transformer hybrid neural network architecture in MutFormer. Details in Supp Fig 3 and Methods. C: Number of channels. K: Convolutional kernel width. H: Number of attention heads. **(c)** Evaluation of model performance on independent test data (mutations on held-out cohorts), across the MutFormer models for each of the 41 SBS mutational signatures. The scatterplot correlates prediction accuracy (as AUC) against the sparsity of the SBS signature’s trinucleotide spectrum (as Gini index, lower for more “featureless” signatures; r = 0.51, p = 0.001). Shaded regions are prediction intervals of the fit. **(d)** Predictive gain, expressed as log2(odds Ratio) and 95% CI over a baseline of random prediction of mutated sites, by MutFormer models for each SBS signature evaluated on independent test data (held out chromosomes).

### MutFormer identifies DNA 21-mer context effect on mutation rate for individual processes

We trained a model using the MutFormer architecture, separately for each mutational signature, including 41 SBS signatures, with between ~17.5M mutations (SBS5) and ~10K mutations (SBS16) in our training set. We evaluated accuracy in predicting mutation rate by the area under the ROC curve (AUC) statistic on a held-out test set of mutations from an independent cohort (mainly ICGC or Hartwig, the cohort with lower number of mutations was considered as the test set). The AUC values range from 0.54 to 0.86 (Fig 1c), suggesting that all signatures were learned by MutFormer better than baseline. Notably, trinucleotide context was not used for training, and every mutation was matched to a nearby non-mutated site with the same trinucleotide context (see Methods, Fig 1b). Therefore these results suggest that the extended DNA context predicts mutation risk for many mutational processes, sometimes with notable accuracy. The accuracies exceeding AUC>0.75 for several types of signatures can be considered high, given that the training data contains only the binary label of if a site is mutated or not. These binary labels are a noisy proxy for the true quantity of interest (quantitative mutation risk), and this noise in training and testing data causes the accuracy estimates to be biased conservatively. The enrichment over random-guessing by the majority of the MutFormer models is >2-fold, and for about a third of the models approximately 4-fold or higher (Fig 1d). Interestingly, signatures with sparse trinucleotide spectra (test set AUC for SBS10a=0.80, SBS10b=0.79, SBS17a=0.79, SBS17b=0.86, SBS7a=0.77) have generally higher AUC values than the “flat” signatures with featureless trinucleotide spectra (test AUC for SBS3=0.56, SBS5=0.55, SBS8=0.59) (Fig 1c, Gini index in the trinucleotide spectrum correlates with MutFormer prediction AUC at r=0.51, p=0.001). This indicates that if a mutational signature is predictable from the trinucleotide, it is further more accurately predictable from the extended DNA sequence context.

Next, we asked what is the width of the extended DNA context that determines the local mutation risk for each mutagenic process. We thus interrogated the influence of each location in the DNA context on mutation rate by *in silico* mutagenesis (ISM), where a certain nucleotide is perturbed, and the difference in the MutFormer predictions are quantified as a measure of contribution of that nucleotide to mutation risk (Fig 2e). We tested statistical significance of the effect of each nucleotide to the risk in the 60 central positions of each 100-mer, contrasting against a baseline of the more distal regions within the 100-mer ([−50bp,-30bp] & [30bp,50bp]), which are expected to have low effects on the mutation risk of the central nucleotide. The statistical test showed significance (p<5.9e-11; t-test, after FDR correction) for all positions tested in [−10,10] window for all considered mutational signatures, suggesting that MutFormer models consistently capture above-random effects of the 20 neighboring nucleotides on mutation risk of the central nucleotide. To prioritize meaningful positions, we used an effect size threshold to highlight nucleotide positions with Cohen’s *d*>1 (Methods). This approach shows that extended mutation context, typically up to 10 nucleotides upstream and downstream, influence mutation rate for many signatures. In particular, the width of the meaningful-effect window of extended DNA context (Cohen’s *d*>1) is on median 15 nucleotides across all the mutational signatures tested (Fig 2e), and is 33, 29, 26 and 24 in signatures SBS44, SBS5, SBS4 and SBS7a respectively. Therefore, DNA elements modulating mutation rates on the genomic “mesoscale”, reported previously for individual example mechanisms, for instance in references^15,8,54^, are in fact common across many mutational processes.

**Figure 2.**
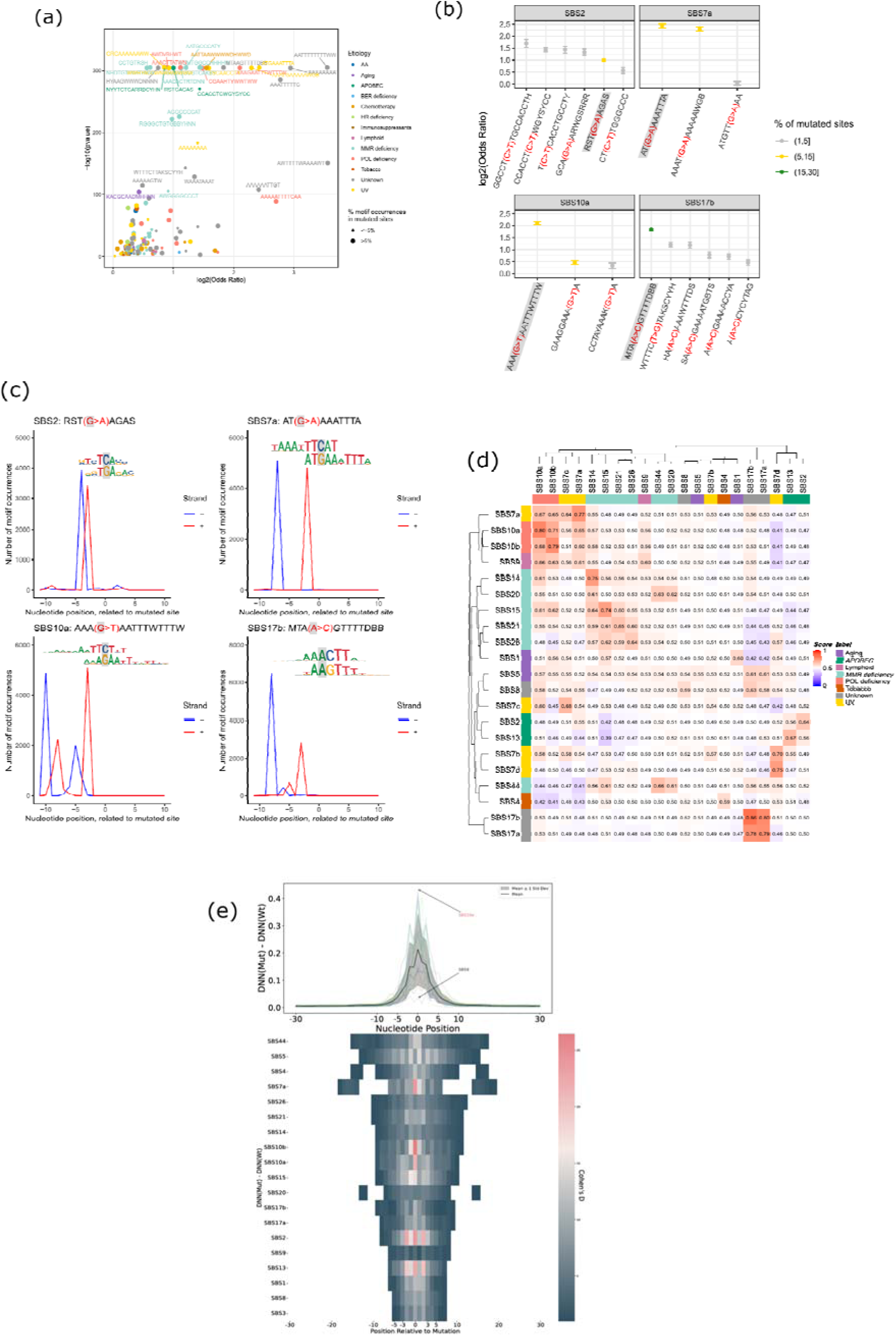
Extended sequence width and motifs, and shared risk etiologies of MutFormers across mutational signatures. **(a)** Extended DNA sequence motifs associated with higher local mutation risk prioritized by MutFormers, for various SBS, identified via STREME analysis by enrichment in high-prioritized oligomers (shown as log2(odds ratio) and corresponding pvalue). **(b)** Effect sizes and 95% CI of the most enriched motifs for SBS2, SBS7a, SBS10a, and SBS17b mutational signatures. Mutated nucleotide in parenthesis and in red. **(c)** Distribution profile showing the local positioning of the four representative motifs relative to the mutated nucleotide. Lines display the number of motif occurrences at each position across the flanking nucleotide windows. **(d)** Asymmetric cross-prediction heatmap displaying pairwise accuracy values (as AUC) when a MutFormer model trained on a specific SBS signature (vertical axis, trainset SBS) is evaluated on a test dataset from the same or another SBS signature (horizontal axis, testset SBS). Clusters of high cross-prediction accuracy reveal shared flanking sequence risk rules, typical among signatures of related etiology. **(e)** Mapping the width of genomic “mesoscale” of sequence dependency of mutation risk, by in silico mutagenesis (ISM) on MutFormers for various SBS signatures. The heatmap quantifies the contribution of individual nucleotide positions (Cohen’s *d* effect sizes, via ISM) within a central 60 bp window across 41 mutational signatures, ordered by the length of the high-effect window (spanning positions with Cohen’s *d* > 1); across SBS signatures, median width = 15 bp. In top panel: DNN, deep neural network (i.e. MutFormer) prediction score; y axis shows subtraction of ISM-mutant and wild-type for every position for individual SBS, and the mean +- s.d. across all SBS.

### Extended DNA motifs associated with local mutation risk gleaned from the neural net

We aimed to interpret MutFormer learned features, by identifying enriched DNA motifs in the highly-ranked predictions (i.e. high mutation risk oligonucleotides) from the various MutFormer models of interest. For that purpose, we sorted a random sample of both mutated and non-mutated sequences from the human genome by their MutFormer mutation risk predictions. Then, we used STREME for motif enrichment analysis^58^ by comparing the top 100,000 predicted sequences against the bottom 100,000 predicted sequences. This yielded, for example for the MutFormer model for the UV signature SBS7a, the 10-mer AT(G>A)AAATTTA motif (Fig 2b,c) (6.27% of the sites top-scored by MutFormer_SBS7a, compared to 1.17% of the sites bottom-scored, OR=5.6). Therefore this DNA motif is one of the top DNA motifs that the SBS7a MutFormer model classifies as mutation risk-conferring; of note various other enriched (i.e. high-risk) DNA motifs were identified for the UV signature and other signatures (Fig 2b). The majority of significantly enriched motifs in the neural net-top hits had more subtle effect sizes (log2 OR<1.0) but were numerous (Fig 2a). This implies that modulation of mutation risk by the local DNA sequence is a distributed property extending genome-wide, rather than being confined to the few known examples of focal mutation hotspots e.g. in the CTCF/Cohesin binding site motifs or in ETS binding site motif or hairpin-forming DNA^15,8,54^.

Next, we considered the motifs recognized by MutFormer trained on the mutational signature SBS10a, observed in DNA polymerase epsilon (POLE)-proofreading mutant cancers (Fig 2c). The SBS10 is known to have an extended context, wider-than-trinucleotide mutagenic motif^36^. As a top hit by number of sites affected, we identified the 14-mer AAA(<u>G</u>>T)AATTTWTTTW motif in the 14.25% of the sites top-scored by MutFormer_SBS10a, compared to 3.32% of the sites bottom-scored (OR=4.3), which matched with the previously reported^41^ RRR(G>T)RRY motif (R= A or G,Y= C or T). We also highlight the top motif in SBS17b, the 12-mer MTA(A>C)GTTTTDBB (Fig 2b,c) with a reverse-complement VVHAAAAC(T>G)TAK (27.6% of the sites top-scored by MutFormer_SBS17b, compared to 7.7% of the sites bottom-scored, OR=3.5). This agrees with and expands motifs previously reported as enriched in SBS17b, the AAAC<u>T</u>T^3^, and AAC<u>T</u>T^59^ and the pentanucleotide signature AC<u>T</u>TA^33^. Regarding APOBEC mutations, we report the most common motif in SBS2, the 8-mer RST(G>A)AGAS (Fig 2b,c) (equiv. to STCT(C>T)ASY; 9.5% of the sites top-scored by MutFormer_SBS2, compared to 4.8% of the sites bottom-scored, OR=2.0), matching and extending the APOBEC3A preferred tetranucleotide YT<u>C</u>A^60^. Using these striking examples, we explored how MutFormer positions the mutation risk-associated DNA motifs relative to the mutated nucleotide (Fig 2c). Despite the full MutFormer window being tens of nucleotides wide, in the 4 examples considered (Fig 2c) we find the motifs are positioned predominantly in a single position (for each of the 2 DNA strands), and that the positioning corresponds to the expected coordinate of the mutation (shown as position +1 in Fig 2c). Therefore these DNA motifs, as recognized in MutFormer models, associate with mutation risk at the nucleotide-resolution. Overall, our MutFormer models can recognize diverse DNA motifs associated with mutation risk, recapitulating several known examples of high mutation risk motifs, reassuring the MutFormers are credible also for the remaining motifs and applicable genome-wide.

### Relatedness of mutational processes by the extended DNA sequence conferring mutation risk

We further asked if some mutational signatures share underlying DNA patterns conferring mutation risk. To assess this, we tested the ability of MutFormer models trained on one SBS signature to predict the other SBS signatures (cross-prediction accuracy). To quantify this, we evaluated pairwise accuracies of a MutFormer model trained on one signature and tested on another signature, expressed as AUC (heatmap in Fig 2d). We observed similar AUC values for cross-predictions between signatures of the same etiology, for example for the SBS7a/b/c/d (Fig 2d) (all four UV-associated) had intra-signature AUCs 0.57-0.77, while the AUCs across signatures SBS7a and 7c (0.64), as well as SBS7b and 7d (0.70). Similar cases were the SBS10a/b (Fig 2d), resulting from DNA polymerase epsilon proofreading deficiency, where AUCs within signatures were 0.79 and 0.80, and across signatures was 0.71. Similar trends were also observed with the MMR deficiency-associated group SBS15/21/26 (Fig 2d), with AUCs across signatures 0.55-0.62 and the AUCs within signatures were 0.64-0.74; also inter-correlated with this group were the baseline MMR signature SBS44 as well as the mutant-polymerase plus MMR signatures SBS14/20 (Fig 2d). The SBS17a/b signature group (Fig 2d), possibly resulting from oxidative damage to free nucleotide pool and/or misincorporation of uracil into DNA, we also saw AUCs SBS17a=0.79, SBS17b=0.86 and the cross-signature SBS17a/b_AUC_=0.78. The two APOBEC signatures, SBS2 and SBS13 were also related by this criterion (Fig 2d), implying some overlap in the flanking DNA sequence motifs that confer the mutation risk to SBS2 and 13. Interestingly, the model for predicting risk of SBS9 -- a lymphocyte-associated signature related with the somatic hypermutation process -- cross-predicts with SBS10a/b and SBS7a/c (Fig. 2d) suggesting that some types of repetitive, A/T rich motifs (see above) might be vulnerable to diverse processes. As another example of this, the putatively NER-deficiency associated SBS8, as well as the universal background signature SBS5, seem to modestly cross-predict DNA sequences with SBS17a/b mutational risk model (Fig 2d). Overall, the signatures generated by related mechanisms are predicted from similar patterns in neighboring DNA, as captured by MutFormer, supporting that different MutFormer models capture the biological underpinnings of each SBS signature.

### Modelling mutation rate at nucleotide-level using MutFormer risk predictions and covariates

Our aim is to systematically analyze recurrent non-coding hotspot mutations for driver potential. Prior work on noncoding mutations in cancer largely aggregated noncoding mutations by region instead of considering individual hotspots, had limited-size WGS datasets thus was not able to measure recurrence nor replication across cohorts well, and most importantly had baseline mutation rate models at the scale of a kilobase^4^ or 10kb^5^ thus did not capture mutation risk due to oligonucleotide effects. The opportunity is provided by the MutFormer, our neural net model to predict background mutation risk from up to 100-mer nucleotide sequences, separately for different mutational processes (which are defined via standard COSMIC signatures). As predictive features, we incorporate these MutFormer local risk predictions for all 41 SBS modelled signatures, jointly with total mutation burden (of tumors bearing the hotspot) and additional covariates (gene expression, replication time; see Methods) to predict the expected number of mutations under neutrality, for each coding and noncoding hotspot with >=2 mutations in a given cancer type.

For the subsequent analysis of significant recurrence in individual hotspots, we expanded our curated WGS data set collection to ~18000 WGS, and for all cancer types at least 2 independent cohorts are available, and for colorectal, ovary, lymphoma at least 3 cohorts (Supp Table 1). Thereby, independent replication of an observed recurrent hotspot can be enforced (i.e. we ignore hotspots occurring in one cohort only, thus lessening impact). For instance, we have 1848 breast WGS, 349 colorectal-MSI WGS, 2144 colorectal-MSS WGS, 1089 ovarian cancer WGS and 1956 kidney cancer samples with WGS. In these five cancer types, considering hotspots with >=2 mutations & replicated across more than 1 cohort, we fit accurately the predicted number of mutations versus observed number of mutations (excluding mutations affecting known driver genes), with Pearson r=0.806-0.862 using a truncated Poisson regression (Methods). This suggests that our background model is broadly accurate, and also that there are large numbers of highly-mutated hotspots that are evolving neutrally (breast, colorectal MSI and MSS, and ovarian in Fig 3a-d, kidney and other cancer types in Supp Fig 6) and drive the correlation between predicted and observed number of mutations. Of note, the data for deriving this hotspot-recurrence neutral model is practically independent of the training data for the MutFormer neural net, which distinguishes unmutated (n=0) from mutated sites (n>=1 recurrences, where the vast majority is n=1), thereby preventing circularity.

**Figure 3.**
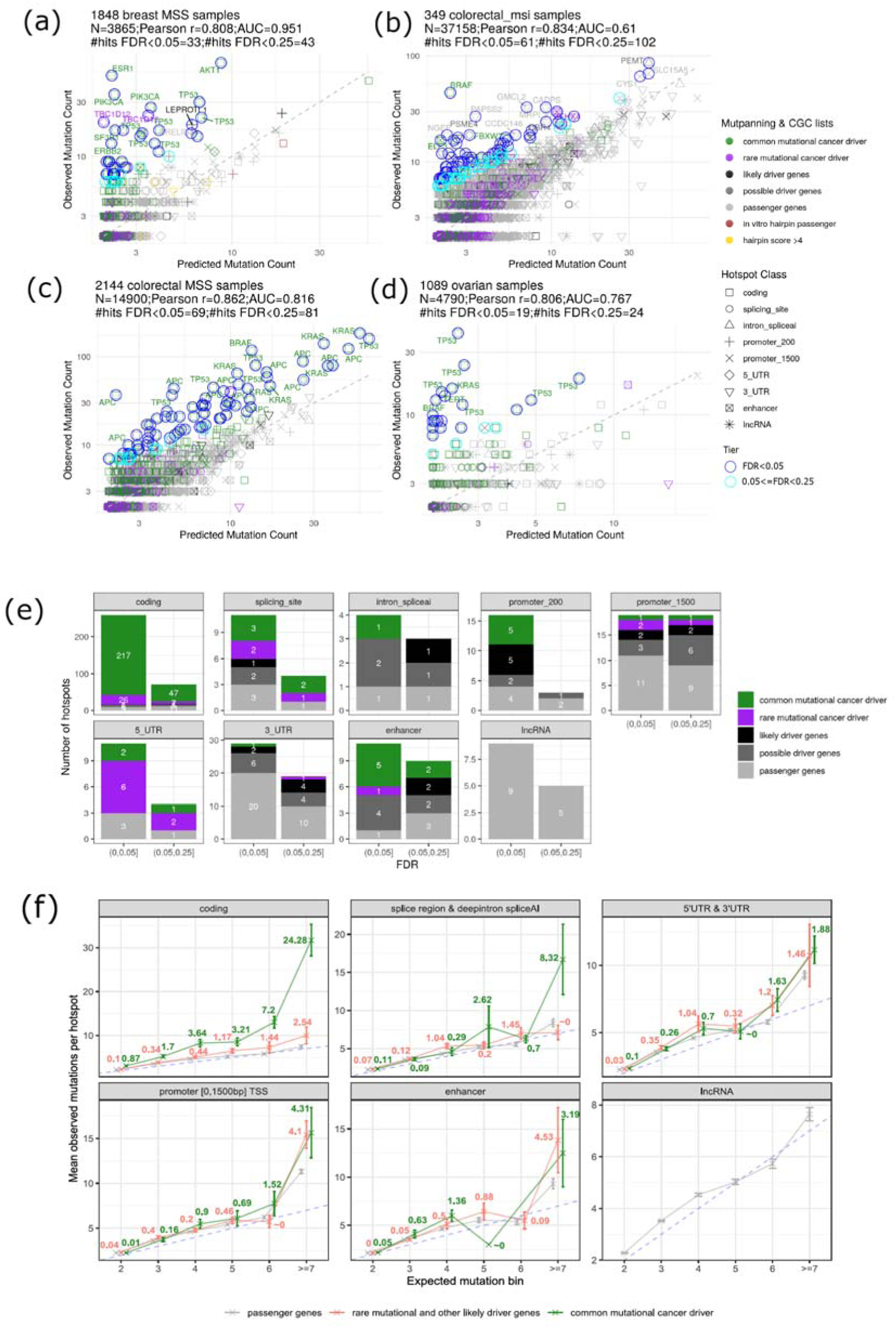
MutFormer-based calibration of background mutation risk and quantification of positive selection across hotspots. **(a–d)** Agreement between predicted (expected under neutrality) and observed somatic mutation counts per individual hotspot across representative multi-cohort datasets: (a) breast (1,848 WGS), (b) colorectal MSI (349), (c) colorectal MSS (2,144), and (d) ovarian cancer (1,089). N denotes the number of hotspots. Other cancer types are in Supp Fig 6. The high Pearson correlation coefficients (r = 0.81–0.86; excluding cancer genes) demonstrate accurate calibration under neutral evolution. Individual points represent unique hotspots, color-coded by driver gene annotation from MutPanning and Cancer Gene Census (CGC) lists (common drivers i.e. known in >=2 cancer types; rare drivers); and from additional diverse sources (see Methods; likely drivers; possible drivers); passenger genes, with shapes indicating specific functional genomic elements (“Hotspot Class”). **(e)** Total number of significant genomic hotspots, pooled across all cancer types (excl. skin cancer) stratified by functional category (coding, canonical splicing site, intron splice region, promoter 200 bp, promoter 1,500 bp, 5′ UTR, 3′ UTR, enhancer, and lncRNA) and partitioned into FDR bins. Counts are hotspot-cancer type pairs. Enrichments over passenger genes are in Supplementary Fig. 9. **(f)** Estimating aggregate numbers of selected mutations, across medium- and low recurrence hotspots, grouped into expected mutation count bins (x axis). The y-axis denotes the mean observed mutations per hotspot, while printed numbers show excess mutation after subtracting the empirical baseline (signal from passenger genes) to quantify positive selection. Dashed line is x=y diagonal. Trends are plotted separately for coding regions, splice regions (incl. deep intronic SpliceAI-supported sites), UTRs, promoters, enhancers, and lncRNA categories. Error bars are standard errors of the mean.

### Various selected hotspots identified, and greatly larger numbers of hotspots lacking signatures of selection

Neutral hotspots were previously reported earlier for various UV-mutagenized promoter hotspots in skin, APOBEC hotspots in DNA hairpins in bladder/breast/lung cancers, CTCF-binding site hotspots in colon/gastric/esophageal cancers, indel hotspots near tissue-specific genes in lung/liver/thyroid/stomach, and AID (somatic hypermutation) hotspots in lymphocytes^9,15,18,8,14,61^. Our analysis generalizes this principle of predicting local mutation risk from various DNA sequence features (and additionally, epigenomic features, see Methods) to many mutational processes and to any location in the genome.

In our comprehensive collection of tissues and signatures, we report 32, 292, 134 and 17 non-coding hotspots with >=10 mutations in the breast, colorectal-MSI, colorectal-MSS and ovarian cancer WGS, respectively; out of these non-coding hotspots, 37%, 70%, 64% and 35%, respectively, have an expected number of mutations >=8 by our MutFormer-based background model meaning, for many of the apparently highly-recurrent hotspots, there is not compelling evidence they are selected and each should be carefully tested. We use these Poisson regression fits to the hotspots in presumably neutrally-evolving sites (meaning, excluding known driver genes and their promoters, enhancers, UTRs and splicing regions) to assess the statistical significance of the residual for each hotspot i.e. the excess mutation recurrence over expectation (see Methods). This test infers selection on the hotspot, while accounting for the local mutation risk and the diversity of molecular mechanisms that could generate that hotspot.

For the majority of the cancer types considered, we identified upwards of 200 hotspots (Supp Fig 7a,b) that had n>=3 mutational occurrences and were thus eligible for testing (Supp Table 2). The numbers of significantly selected hotspots for a cancer type were typically in the tens, including coding regions (Supp Fig 7c), even at permissive FDR thresholds. In particular, no hotspots with n<=4 mutations reached significance at FDR 25%, with rare examples in liver, pancreas and uterus MSS with FDR<50%; only two dozen hotspots at n=5 were FDR 25% significant, tallying across all cancer types (Supp Fig 7d). This hints that the testing framework is conservative; we assess our approach further below, using for benchmarking the better-studied cases of coding region hotspots, to which our method is equally applicable as to the noncoding ones.

### Analyses of hotspots in coding regions suggest a conservative test for hotspot selection

As a control of the methodology, we firstly consider specifically the coding region mutational hotspots. Driver genes activated/inactivated by coding region mutations have been well studied and thoroughly catalogued^62^. Moreover in coding regions there exist good estimates of variant impact: synonymous mutations are in the majority of cases neutral, and missense mutations can be sorted into low-impact and high-impact accurately by predictors such as AlphaMissense^63^, shown to be applicable to cancer driver mutations^64^. This makes it possible to infer selection in a manner orthogonal to recurrence, and provides a way to test the reliability of our MutFormer-based methodology for detecting positive selection on individual hotspots.

We tested n=75744 coding region hotspots across 18 cancer types in total; note that if the same hotspot occurs in multiple cancer types, they count as separate instances, meaning we do not perform a pan-cancer analysis here, but simply consider the analysis of the individual cancer types in aggregate (shown in Fig 4a). Each hotspot had to be observed across >1 cohort. Of these, we considered for statistical testing the n=11262 hotspots with n>=3 mutational occurrences, and where the direction of the effect was towards enrichment. Upon a Benjamini-Hochberg adjustment for multiple testing, 277 coding region hotspots were significant at FDR<5%, 363 at 25% and 455 at 50%.

**Figure 4.**
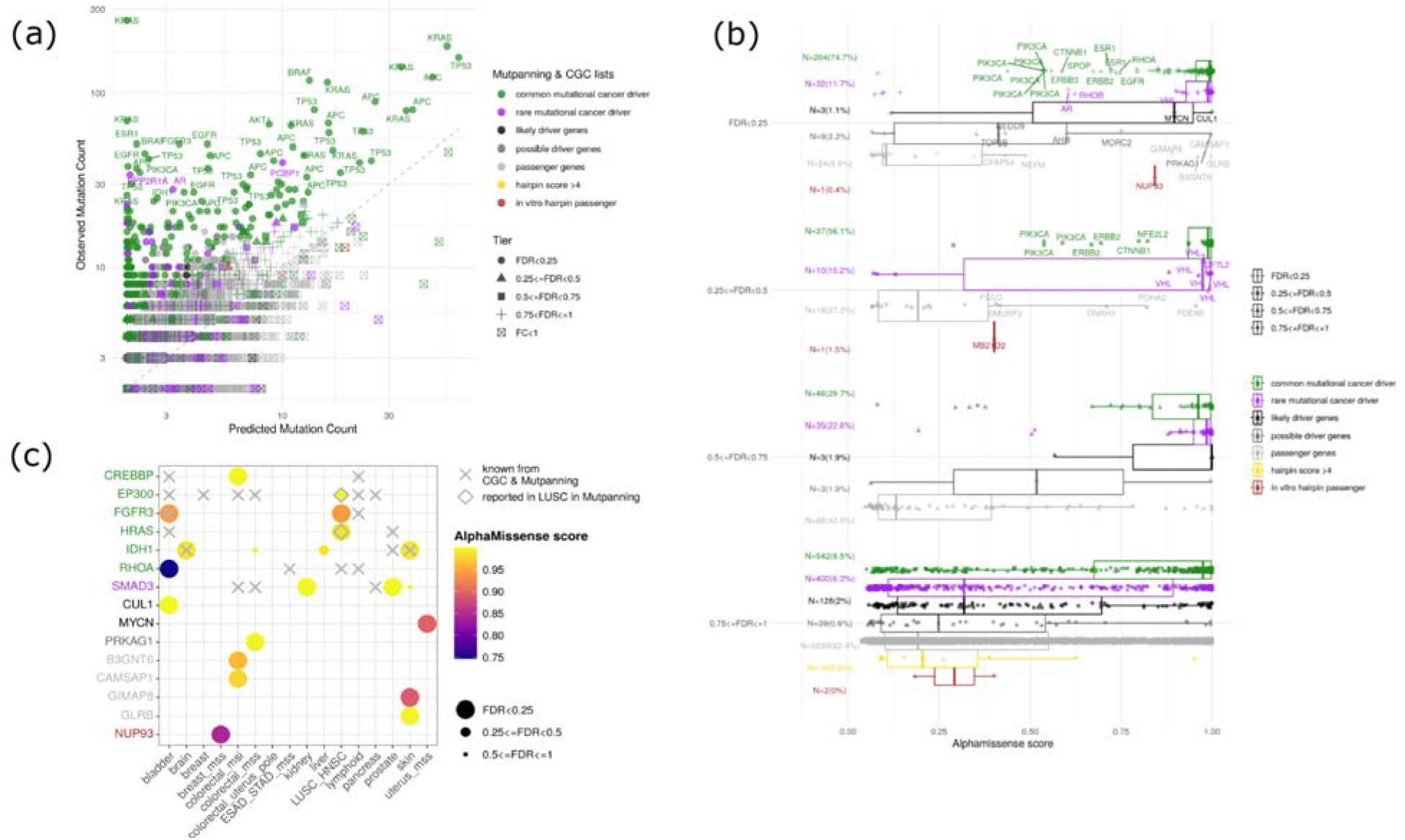
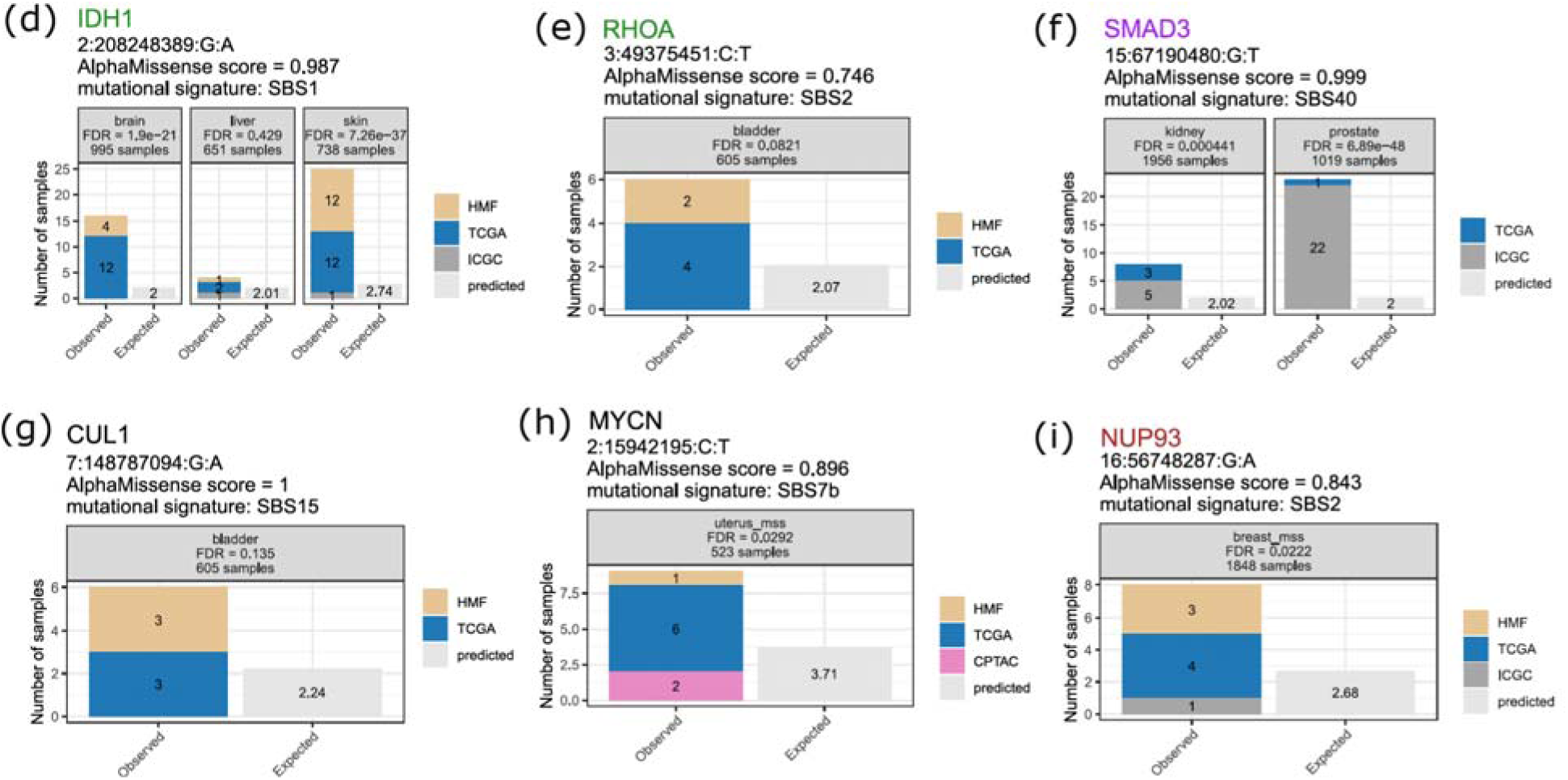
Validation of the hotspot-recurrence methodology using gene coding regions, and identifying rare tissue/driver combinations. **(a)** Observed versus predicted mutation counts for 75,744 coding region hotspots across 18 cancer types. Points represent individual hotspots in a particular cancer type, colored by driver category (as in Fig 3), with shapes indicating FDR tiers. **(b)** Orthogonal calibration and empirical FDR assessment of the methodology using AlphaMissense pathogenicity scores in coding regions. Distribution profiles are stratified across nominal FDR significance tiers and split by driver versus passenger status, confirming a low empirical FDR (9.6% for the group with nominal FDR<25%), and a high truepositive density among suggestive hits with 25%<=FDR<50%. Categories “hairpin score >4” and “in vitro hairpin passengers” pertain only to APOBEC signature hotspots (SBS2, SBS13). **(c)** Tissue-specificity matrix displaying significant coding-region hotspots across 18 tumor types. Dots are colored by AlphaMissense scores and sized by statistical significance tiers. **(d–i)** Detailed breakdown of example coding hotspots in genes, for cancer types where they are known drivers (marked with X in panel **c**) and additional cancer types where they are identified as significant or marginal in our analysis. Bars contrast the observed mutation counts (stratified by data source cohort) against the predicted neutral baseline count.

We considered the reliability of our significance estimates, in light of calibration of p-values against a theoretical distribution, the enrichment in cancer genes and enrichment in high AlphaMissense scores. We interpret that the p-values (and thereby, FDRs) from our tests to be unbiased or conservatively biased, based on inspection of quantile-quantile plots of p-values for various cancer types. In particular, we observed that 13 out of 18 cancer types have well-calibrated or deflated p-values (lambda <=1.0, Supp Fig 8; shown for hotspots in all genome regions including coding and various noncoding regions), while the remaining 5 cancer types had mild inflation (1<lambda <= 1.32 (uterus MSS)). Considering the FDR<25% threshold, the enrichment in known cancer genes and the estimates of pathogenicity by AlphaMissense score suggest our FDR estimates are conservative. Firstly, of the significant coding hotspots at nominal FDR<25% that also had AlphaMissense score available, 204+32 are in known driver genes (common + rare driver genes, respectively), while only 24+1 are in passenger genes (Fig 4b). This would imply an empirical FDR (eFDR) of 9.6% for this FDR<25% hotspot group, under the conservative assumption that our set of passenger genes do not contain selected hotspots. By same criteria, the coding hotspots significant in the marginal FDR 25-50% range (Fig 4b) would have an eFDR=28.7%, suggesting that even this FDR<50% group has utility for driver hotspot discovery. Secondly, the AlphaMissense score distributions in the 204+32+24+1 driver and passenger gene hotspots at FDR<25% suggest only 26 hotspots with AlphaMissense<0.5 implying 9.9% eFDR by this criterion; the hotspots with nominal FDR 25-50% would by same criteria have eFDR=30.3%, both estimates very close to eFDRs obtained via driver gene enrichment criteria. Based on these tests conducted on coding regions, henceforth we consider hotspots at FDR<25% significant, and those at FDR 25-50% can be considered suggestive hits, where approximately two out of three hits in this FDR 25-50% category are likely to be true positives.

### Tissue-specificity spectrum of cancer driver genes refined through coding hotspot analysis with local mutation risk model

Overall, in gene coding regions we identified 330 significantly recurrent hotspots at FDR<25%, of which 259 have FDR<5% (not considering AlphaMissense score, Fig 3c), here counts implying unique locus-cancer type combinations. Of those hotspots, 264 are in common cancer driver genes (previously reported mutational drivers in multiple cancer types, according to MutPanning significant enrichment and/or Cancer Gene Census catalogs), 33 in known rare driver genes (previously reported mutational drivers in one cancer type in same catalogs), 13 in likely driver genes or possible driver genes (including dosage-sensitive genes from Cancer Gene Census, from SWAN analysis of copy number alterations, and additional genes we identified in bespoke search), and 20 in presumably passenger genes.

We reasoned that our analysis of gene coding driver mutations, which is hotspot-focussed, draws on a large sample size and is boosted by AlphaMissense as independent support, may improve over previous analyses in identifying additional cancer types where a driver gene is selected (Fig 4c). For instance, we report high confidence hotspot hits for *SMAD3* in kidney cancer and prostate cancer, which have ~4-20 fold recurrence enrichment over expectation (Fig 4d); this complements existing annotations via CGC or MutPanning that implicate *SMAD3* in colorectal cancer and pancreas cancer (Fig 4c). The *IDH1* oncogene is a typical driver for glioma, and our analysis further supports *IDH1* driver roles in melanoma (proposed through smaller-scale analyses^65,66^ and in liver cancers, with ~2-9 fold enrichments suggesting positive selection, as well as AlphaMissense support (Fig 4e). With similar confidence, we report selected occurrences of other known driver genes in additional tumor types (Fig 3c), in particular *CREBBP* driver hotspots in colorectal-MSI cancers and *FGFR3* hotspots in the (here jointly considered) lung squamous and head-and-neck squamous cell cancers.

Next, we consider selected hotspots in coding regions of genes that are not listed as mutational drivers in the comprehensive catalogs of CGC and MutPanning, but have other evidence of involvement in cancer (here, listed as “likely driver genes” or “possible driver genes” categories, commonly with involvement via gene copy number changes, and/or via a literature search, see Methods). For example, we identify *MYCN* hotspots in uterus-MSS cancer (Fig 4g), *CUL1* hotspots in bladder cancer (Fig 4h), RHOA hotspots in bladder cancer (Fig 4i) and the *NUP93* hotspots in breast cancer (Fig 4f) that are all significant at FDR<25%, have >2-fold enrichment over expectation and recur across multiple cohorts, and have high AlphaMissense scores of >0.8; *NUP83* has a caveat of prior report of hypermutable hotspot by APOBEC in a DNA hairpin^15^. Meeting the same criteria for selection and driver impact, we also list examples of apparent passenger genes that nonetheless bear significant hotspots with AlphaMissense>0.8, including *GIMAP8*, *CAMSAP1*, *GLRB*, and *B3GNT6* (Fig 4b, Fig 4c).

### Selected noncoding hotspots are very rare, however strongly enriched at known cancer genes

While coding regions of cancer genes have been studied extensively in various large scale analyses including that of exome sequencing and panel sequencing, the prior analysis of non-coding regions has typically been constrained by lower availability of WGS. Analyses ranged from hundreds of tumors with WGS^67,17,55,68^ to up to approximately 2.5 thousand WGS in the PCAWG cohort^2,9,69,42,5,70^, with a more recent analysis combining cohorts to reach 3.9k WGS to search non-coding drivers^4^. Our combined dataset has 18,317 WGS tumor samples -- a ~7-fold increase over PCAWG size -- greatly improving sensitivity. Taking advantage of the larger sample size, our statistical methodology, uniquely for non-coding analyses, tests significance of each individual hotspot rather than pooling mutations across nearby loci, potentially enabling more precise identification, and facilitating removal of artefacts. First, we turn towards selected promoter mutations, where one example is very well established (telomerase gene *TERT*^55,71^), and two other examples have moderate evidence of activity (promoters of *FOXA1* and *CDC20* genes^72,73^, and we asked if our comprehensive analysis could reveal additional driver hotspots in promoters.

Here, promoters were defined as 200 nt upstream of TSS (Pr200), where we have tested for significance 3775 hotspot-cancer type pairs (henceforth, “hotspots”) with n>=3 observed mutations (requiring n>=4 in extremely mutated cancer types, colorectal_MSI and colorectal+uterus_POLE) and with non-negative effect size (log-enrichment), meaning, not depleted in observed over expected mutation counts (the below-diagonal points on plots Fig 3a-d or Supp Fig 6). Of note, these noncoding hotspot analyses span various cancer types, but from the counts reported here we exclude skin cancers (whose various significant hits are shown in Supp Fig 6), which should be considered with more caution given the extreme local mutagenesis due to UV damage at ETS binding sites^8,14^. We identified 19 significant TSS hotspots at FDR<25% (Fig. 3e), of which 5 in known driver genes, and additional 5 in likely driver genes. The expectation based on total numbers of driver and passenger genes implies a ~3-fold enrichment over random, for various cancer gene groups considered (albeit with broad confidence intervals, see Supp Fig 9). Second, we consider a more extended promoter-proximal region of 1500 nt to 200 nt upstream of gene TSS (Pr1500), with 5854 tested hotspots and 38 significant hotspots. Of those, 5 were in known driver genes and 4 additional significant hotspots in likely driver genes (Fig 3e), implying a ~3-10 fold enrichment over a random distribution across genes (shown separately for groups of known common, known rare, and likely driver genes, Supp Fig 9).

In addition to promoters, we further considered hotspots in enhancers, gene untranslated regions (UTRs), splice regions including deep intronic, and non-coding RNA genes. Considering TSS-distal enhancer elements from a catalog generated from cancer cell lines and genes inferred to be linked to these enhancers^74^, we tested 2205 to identify 20 significant hotspots, of which 10 in known cancer driver genes (Fig 3e) and 2 in likely driver genes, thus >=10-fold enriched with respect to random gene distributions across categories (Supp Fig 9). Considering UTR hotspots, in 9799 tests we observe 63 significant hits (tally shown separately for 5’ UTR and 3’ UTR in Fig 3e), 13 in known cancer genes, and 6 in likely cancer genes, implying ~3-30x concentration of selected UTR hotspots in cancer driver gene groups (Supp Fig 9). Next, as a noncoding category we considered splice sites and deep intronic mutations in splicing-sensitive loci^75^ (SpliceAI>0.10; shown separately in Fig 3e), where out of 3713 tests we called 22 significant hotspots, of which 9 in known driver genes and 2 in likely driver genes, thus ~10-50-fold enrichment (Supp Fig 9). Finally, out of 7460 tested, we report 14 significant hotspots in lncRNA genes (Fig 3e). Taken together, these results suggest that while cancer genomes contain a large number of noncoding hotspots with medium or high recurrence, for the vast majority of them there is not robust evidence of positive selection. The few cases presenting signatures of selection show strong enrichments in known mutational-driver cancer genes and other genes with driver potential (such as identified dosage-sensitive cancer genes and genes with literature support of cancer roles). These cases of selected hotspots that are in noncoding elements proximal to driver genes or suspected cancer genes are plausible candidates for further assessment using variant effect predictor tools such as AlphaGenome^76^, which we perform below.

### Modest but pervasive selection on medium-recurrence, abundant hotspot mutations

Next, we reasoned that the apparent extreme rarity of selection on non-coding hotspots is likely to result from limiting statistical power of the dataset to detect selection in less recurrent hotspots. Such lower-recurrence hotspots -- here operationally defined as having n=2-3 mutations in a cancer type -- and medium-recurrence hotspots with n=4-6, are very numerous and thus even a small fraction of selected non-coding mutations amongst them could make a notable contribution towards the driver tally of tumor genomes. In support of this notion, in our analysis of hotspots in coding regions, we noted that many nonsignificant hotspots, with FDR>75%, were nonetheless possible drivers, based on them being enriched in known cancer genes and based on their commonly high AlphaMissense scores (Fig 4b, bottom panels). Thus, despite reporting a comprehensive WGS dataset, our analysis is likely underpowered for the low-recurrence hotspots. For example, in our assembled set of 2144 colorectal MSS cancers, none of the hotspots recurrent in n<=6 samples (i.e. 0.27% of the cohort) were FDR<25% significant (Fig 3c; Supp Fig 7c); this includes the 5440 hotspots with n<=6 that were in coding regions (Fig 3c). In 1089 ovarian cancer WGS, only 2 out of 1574 coding region hotspots with n<=6 (0.55%) were FDR<25% significant (Fig 3d, Supp Fig 7c), and in 1848 breast cancers, only 6 out of 913 coding hotspots with n<=6 recurrences (0.32%) were significant (Fig 3a, Supp Fig 7c). Based on this, we infer our analysis is generally underpowered to detect significant hotspots in the sub-0.5% rate of recurrence in a cohort. However we also observe that there are many coding region hotspots in known driver genes (thus, a priori strong candidates for driver hotspots) with n=4 to 6 recurrences: 25 in breast, 114 in colorectal MSS, 62 in colorectal MSI, and 23 in ovary (Fig 3a-d; see other cancer types in Supp Fig 6). This supports that many driver events do exist in that range of sub-0.5% recurrence; this can likely be extrapolated from the case of coding hotspots addressed above, also to the case of non-coding hotspots.

We asked how many selected mutations can be inferred to exist in such medium-recurrence hotspots, estimated as an excess of observed over expected mutations considered collectively over hotspots in a particular category, without testing significance of individual hotspots. We restricted this to driver genes with associated non-coding elements (promoters, enhancer, 5’ UTR, 3’ UTR, splice regions and deep-intronic splice altering regions). We used passenger genes as a baseline in this analyses, and subtracted their signal (mutation excess, encouragingly fairly close to neutral expectation; Fig 3f) from the driver genes mutation excess.

Firstly we considered medium-recurrence coding mutations. At hotspots where n=4, 5 and 6 mutations are expected in a given cancer type, an excess of 3.6, 3.2 and 7.2 mutations are observed in common driver genes (Fig 3f, having subtracted the excess in passenger genes), and excess of 0.4, 1.2 and 1.4 in rare driver genes, respectively (Fig 3f). Thus in these low-recurrence categories of n=4 to 6, roughly half of the hotspot mutations in coding regions of common cancer genes are driver mutations, and roughly one-tenth to one-fifth in rare cancer genes are drivers.

Moving on to noncoding mutations, using the same methodology we address promoter 200 and promoter 1500 regions, considered jointly here. In promoters of common cancer driver genes, the hotspots with n=4, 5 and 6 expected mutations have an estimated excess of 0.9, 0.7 and 1.5 selected mutations, respectively (Fig 3f), and rare cancer genes have excess of 0.2, 0.5 and ~0, respectively (Fig 3f). Therefore approximately one-fifth of medium-recurrence promoter mutations in cancer driver genes may be selected. Considering other noncoding elements --UTRs, splice regions (incl. selected deep intronic parts) and enhancers -- in the medium-recurrence hotspots with n=4-6, there is modest excess of putatively selected mutations in cancer driver genes, albeit lower magnitude than in promoters and considerably less precisely estimated (all in Fig 3f), precluding quantification in this study. As a counterpoint to the above, the noncoding elements (promoter, UTR, splice region and enhancer) hotspots with low recurrence i.e. n=2 mutations in our data show negligible signatures of selection, even in common cancer driver genes (Fig 3f). Coding gene hotspots are however selected even with n=2, consistent with overall considerably weaker selection on noncoding elements than on coding regions. Overall, we conclude that medium-recurrence hotspots in noncoding elements, in the range of 0.1%-0.5% mutated samples in a cohort, may harbor a non-negligible fraction of driver hotspots even though they are not powered for reaching significance in a genome-wide test.

Therefore in the following analysis we perform restricted hypotheses tests, testing only noncoding elements associated with cancer genes (common, rare, likely cancer genes and suspected genes group) thereby lessening the multiple-testing burden and effectively increasing power to detect noncoding drivers in medium-to-low recurrence hotspots.

### Evidence for selected splice-region and deep intronic hotspot mutations in TP53 and VHL

Next, we asked if we can provide orthogonal support to the significant non-coding mutation hits, by employing variant effect predictions from various genomic AI tools, serving *in lieu* of the dN/dS tests employed for coding DNA. We first considered putative splice altering mutations that affect known, or possible cancer genes. Here, in addition to the canonical splice site (2 nt at intron border) where splice-altering effects are presumed for all mutations, we also considered all sites in the broader “splice region” (intronic 3-20 nt next to the splice site). Additionally, we aimed to study all deep-intronic sites (i.e. further than 20nt) in known cancer driver genes where the SpliceAI tool predicted a possible splicing effect of a hotspot to be above baseline. This SpliceAI threshold broadly corresponds to the median of the passenger-genes intronic hotspot effects and is intended to, permissively, enable consideration of various tentative splice effects. Of note, as extra evidence for the examples of significant hotspots, we will require validation by an additional splicing predictor, AlphaGenome.

We observed 28 significant splicing hotspots at MutFormer selection FDR<25%, and 22 more at FDR 25-50% (Fig 5a, 5b). We note 36 more hits at the nonsignificant FDR 50-75%, apparently similarly enriched with strong SpliceAI scores as the better FDR categories (Fig 5b). Across these significantly selected categories, there are common *TP53* splicing hotspots (12 hotspot-tissue pairs at FDR<50% and 8 more at FDR 50-75%) and *VHL* splicing hotspots (10 hotspot-tissue pairs at FDR<50% and 4 more at FDR 50-75% Fig 3b). The remainder of selected hits are provided in Supp Table 3 and visualized in Fig 5c.

**Figure 5.**
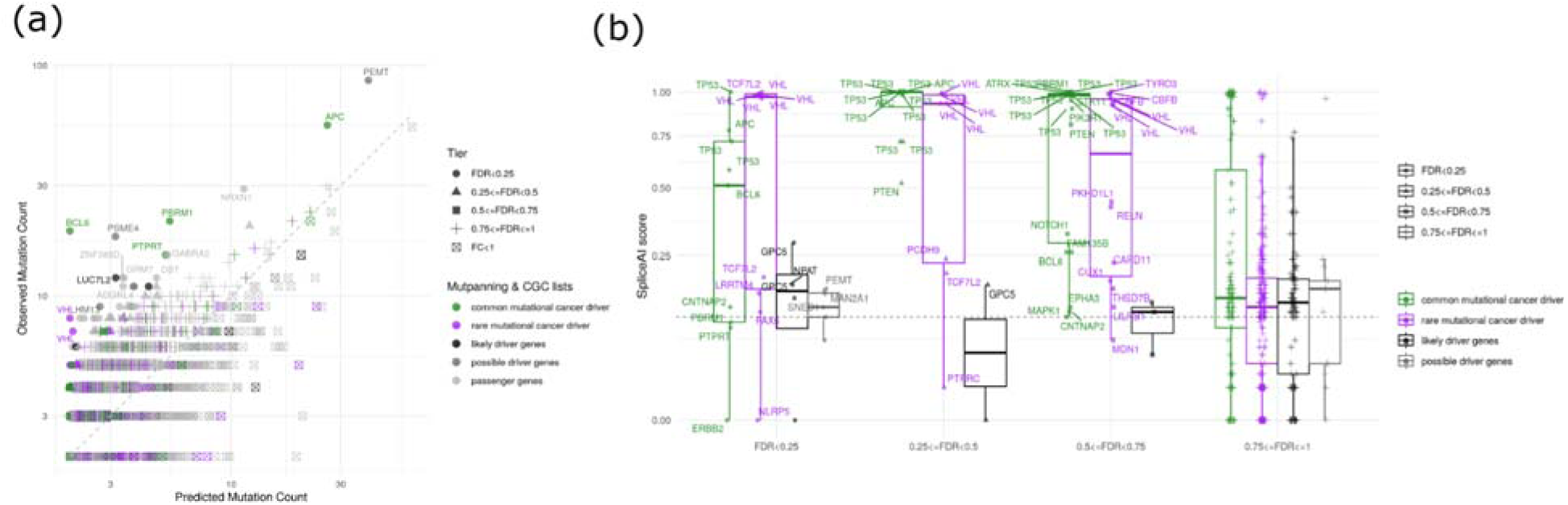
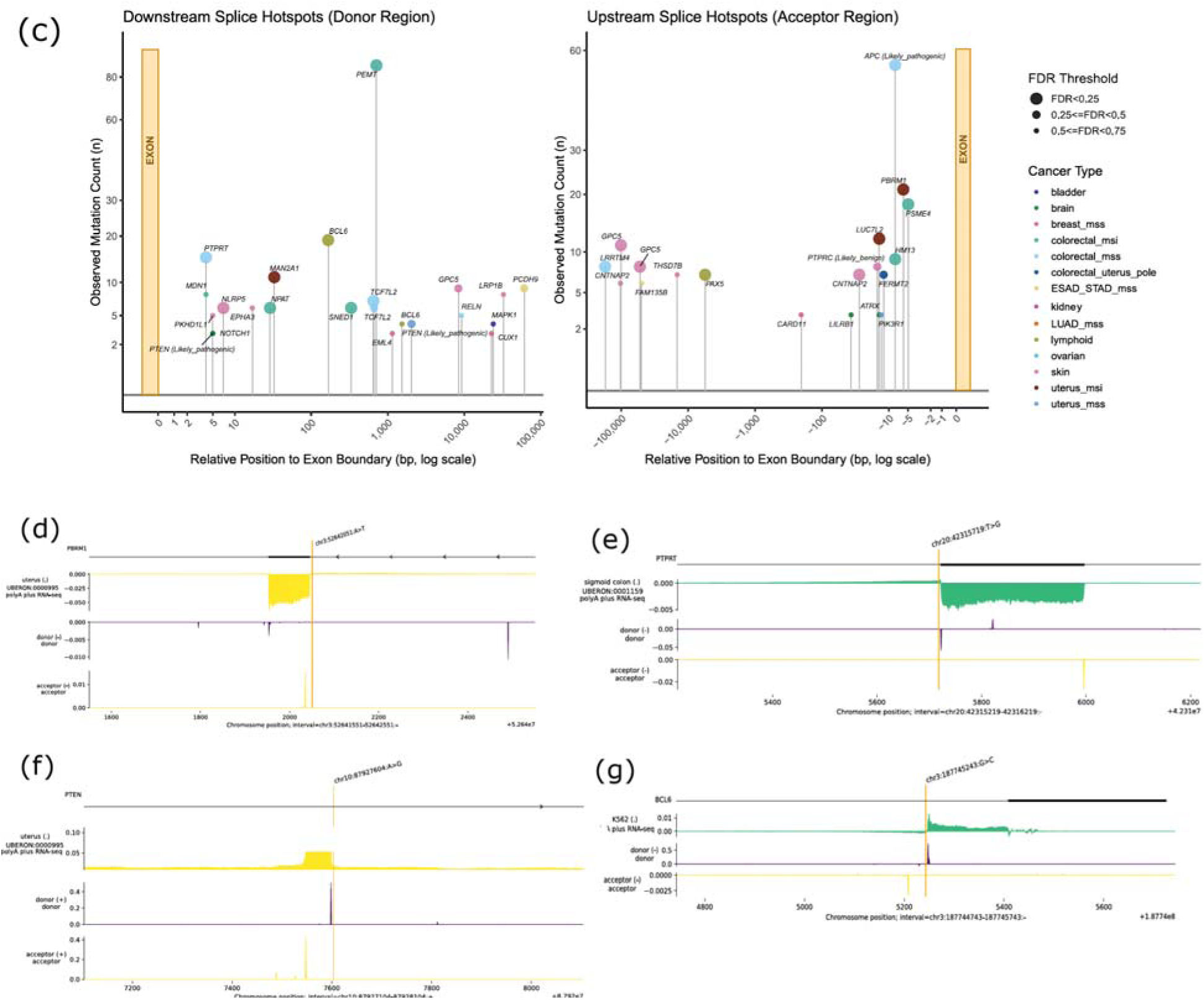
Selection on noncoding splice-region and deep intronic hotspots, additionally supported across AI predictors. **(a)** Observed versus predicted mutation counts across somatic hotspots mapped to canonical splice sites (2 nt in the intron), broader splice regions (as per VEP, 3-8nt in the intron), and deep-intronic sites with possible splice effects (SpliceAI>=0.1). Dots are stratified by gene categories, with shapes mapping to FDR tiers. One point is one hotspot-cancer type combination. **(b)** Distribution of SpliceAI scores across tested hotspots grouped by significance tiers demonstrates a progressive of high-confidence predicted splicing-disruption events (SpliceAI>0.50) in the prioritized selection bins. **(c)** Lollipop-plots showing recurrent somatic mutations flanking exon boundaries in downstream acceptor (left panel) and upstream donor regions (right panel). Relative distances to the nearest exon splice junction (0 bp) are shown on the X-axis using a log scale. Individual points are color-coded by t cancer types, and their sizes reflect FDR tiers. Hotspots present in ClinVar dataset are labelled with their corresponding pathogenic effect. **(d–g)** AlphaGenome validation profiles of selected driver splice-region (**d, e**) and deep-intronic **(f, g)** mutation hotspots across distinct tissue cohorts. Predicted RNA-seq signal difference, and donor/acceptor site strength difference, is shown, aiming to match the cell type with AlphaGenome options as much as possible. **(d) *PBRM1*** splice-region mutation in uterus MSI cohorts (n = 21), demonstrating predicted exon skipping. **(e) *PTPRT*** splice-region mutation in colorectal MSS cancers (n = 15). **(f) *PTEN*** deep-intronic hotspot in uterus MSS tumors (n = 4). **(g) *BCL6*** deep-intronic hotspot in lymphoid malignancies (n = 19), Additional AlphaGenome validations for splice-altering hotspots in genes *TP53*, *VHL*, *BCL6*, *PTEN* and *TCF7L2* are shown in Supp Figs 11, 12, 13, 14.

We found previous reports for 19 out of 20 TP53 splicing hotspots in ClinVar (with 1 remaining TP53 hotspot mentioned as uncertain significance), and 12 out of 14 VHL hotspots, and the 2 remaining were to our knowledge uniquely identified in our study (denoted in Supp Fig 10). A new *TP53* hotspot example that is plausible (although FDR=74% in our test) and reported in ClinVar with uncertain significance is <u>chr17:7,675,989</u>:C>A, which resides at +5 intronic position after TP53 exon 4, and we observe n=4 recurrences in the ovary cancer. This splice-region TP53 hotspot validated in AlphaGenome that indicates that the donor site is abolished and predicts that upstream exon 4 is truncated (Supp Fig 11a). A *VHL* hotspot example not reported in ClinVar and recurrent at n=5 kidney tumors and selected at FDR=16% is <u>chr3:10,142,189:T>A</u>, which is in the canonical splice site +2 downstream of exon 1, and our AlphaGenome validation indeed shows donor site disruption and a predicted extension of the upstream exon 1 (Supp Fig 11b). The second plausible VHL hotspot not reported in ClinVar (at chr3:10,146,639:A>T, spliceAI = 0.92) was detected in our analysis at FDR=63% and n=3 tumors.

### Additional selected hotspots, proximal to splice sites in further cancer genes

Next, we consider splicing effects on genes in addition to *TP53* and *VHL*, where significant hits included at least one hotspot-cancer type pairs with significance (selection FDR<25%) and with non-negligible SpliceAI score: *TCF7L2*, *APC*, *BCL6*, *GPC5*, *LRRTM4*, *CNTNAP2*, *PAX5*, *PBRM1* and *PTPRT*. There are additional such hotspots in the marginal FDR<50% and 75% ranges including in *PTEN* among other genes (Fig 5c); in these FDR bins the distribution of SpliceAI scores was nonetheless considerably shifted from the non-selected hotspots (FDR>75% range) suggesting various valid hits in the FDR<50% range. Using AlphaGenome, we further attempted to validate the splicing effects of various putatively selected hotspots in these genes.

First, we consider the 3 example splicing-altering hotspots in *APC*, which are either known or well anticipated since they lie directly in a canonical splice site. First, a splice-altering hotspot is detected at FDR<25%, 8bp away from exon, with 55 occurrences in colorectal-MSS cancers. This splice-altering somatic mutation is known, and interestingly previously ascribed to SBS88 mutational signature^77^, reported as c.835-8A>G^78^, and additionally known in ClinVar to contribute to familial adenomatous polyposis. In addition to the known mutation, we identify 2 more known *APC* hotspots selected at FDR<50%, both in canonical splice site and both reported in ClinVar as pathogenic in the germline for polyposis/hereditary cancer risk (Supp Table 3).

Next, we consider in the *PBRM1* gene a hotspot 6bp distance from exon boundary (i.e. outside of the canonical splice site) with n=21 in uterus-MSI with FDR=7e-10, that validates in AlphaGenome which predicted exon skipping (Fig 5d). Additionally in *PBRM1* there is a marginal-significance (FDR=63%) hotspot n=3 in kidney in the canonical splice site (at 1bp distance from exon) and consistent with that, having spliceAI=1. Neither PBRM1 hotspots are reported in ClinVar.

Next, in *PTPRT* gene there is a strongly recurrent hit in colorectal-MSS (n=15, FDR=0.0001), proximal to the splice site at 4bp distance from exon, not reported in ClinVar, and which we validated in AlphaGenome predictions as causing exon skipping (Fig 5e). *PTPRT* was reported as a mutational driver gene in a recent large-cohort study of colon cancer WGS^78^, but we do not have knowledge of reports of intronic, splice-altering driver mutations in *PTPRT*.

### Deep intronic predicted splice altering selected hotspots in cancer driver genes

Next, in *BCL6* there is strikingly a deep intronic hotspot (167bp distance from exon), with n=19 mutations in our cohort of lymphoid cancers and FDR=2e-31 and spliceAI=0.51, predicted to generate an alternative splice donor site (Fig 5g with AlphaGenome validation), and not reported in ClinVar. An additional splicing hotspot in BCL6 at marginal significance (n=4 recurrences in lymphoid, FDR=70%, spliceAI=0.26) was noted (Fig 5b), with very deep intronic positioning (1537bp away from exon) and AlphaGenome suggesting a possibility of partial intron retention (Supp Fig 11). We note that the mutations also overlap an antisense mRNA gene and we cannot formally rule out mutation activity via affecting that ncRNA.

Next, the deep-intronic *PTEN* hotspot chr10:87927604:A>G was selected at FDR=38% (Fig 5b, recurrences in n=4 uterus-MSS cancer type, spliceAI=0.52), is situated between *PTEN* exon 3 and exon 4 at 2,407bp distance. This variant was reported likely pathogenic in germline, in ClinVar, via an observation in a hereditary polyposis patient^79^, making plausible also the somatic oncogenicity reported here. We find pathogenicity supported by AlphaGenome predicting exonisation of a part of the intronic sequence (Fig 5f), broadly corresponding to exon 4 of 10 in an atypical *PTEN* transcript PTEN-210 (<u>ENST00000688922.2)</u>, which is an NMD substrate. We note a second putative driver *PTEN* hotspot chr10:87931094:G>A with a marginal FDR=57% but high spliceAI=0.81 (n=3, brain cancer), 5bp away from the exon. The corresponding germline variant is reported to be likely pathogenic in ClinVar for various cancer risk phenotypes including Cowden syndrome and glioma susceptibility, thus plausibly being pathogenic also as somatic mutation. AlphaGenome predicts exon skipping (Supp Fig 13).

Next, in *TCF7L2* gene, encoding a DNA-binding nuclear effector of Wnt/β-catenin signalling, in addition to one hotspot (n=6 colorectal-MSS, FDR<25%) with unsurprising splice effects as it is located in splice site, we also identify an interesting case of two nearby, apparently deep-intronic (650bp and 662bp away from exon in the MANE transcript, TCF7L2-206) hotspots with n=7 and n=6 recurrence in colorectal-MSS at at chr10:113160642:C>T and chr10:113160654:G>A, at FDR<25% and 25%<=FDR<50% respectively. For these 2 somatic mutations, AlphaGenome predicts exon skipping, for an exon observed in two non-MANE transcripts that GTEx reports as expressed in colon (Supp Fig 14, <u>GTEx</u>). Therefore, this is a likely example of exonic coding variants that act via altering splicing of a coding exon^80^, here shown for an alternatively spliced, C-terminal region exon 13 in the *TCF7L2*, where exon skipping in this region are known to alter the protein product and Wnt/β-catenin target promoter transcriptional output^81^. The proximity of the 2 mutational hotspots adds additional evidence of driver potential.

At more moderate SpliceAI thresholds, additional putative splice-altering, hotspots with inferred selection at FDR<25% were seen in *PEMT*, *CNTNAP2*, *LRRTM4*, *GPC5* (two hotspots) and *PAX5*, and additionally *PCDH9* and *GPC5* at FDR<50%. After inspection of AlphaGenome predictions, we noticed that the most recurrent putative splicing hotspot in our dataset (*PEMT*, n=86, colorectal MSI, spliceAI=0.16, 713bp distance to the exon) show modest effect on acceptor track, and low effect in RNA-seq and donor track therefore we do not see compelling support for the splicing effect of *PEMT* intronic hotspot, although we do not rule out selection via other mechanisms. Of the other above-mentioned, moderate SpliceAI hotspots, only two have some AlphaGenome support: a GPC5 hotspot at chr13:91591818:C>T (n=11 recurrences in skin) and PAX5 hotspot at chr9:37026417:G>C (n=7 in lymphoid) predicted an outcome (exonization) without obvious noise in AlphaGenome splicing coverage output, and would therefore be prioritized more highly than the others.

### Transcription start site-proximal hotspot mutations include TERT drivers in 5’ UTR

Of the non-coding somatic mutations, those affecting gene promoters would a priori be expected to result in some driver mutation. The expectation contrasts an apparent dearth of promoter mutations with clear driver potential, where *TERT* is the only incontrovertible example currently. As in the analyses above, we consider only TSS-proximal regions of the genes with a known, likely or possible driver role, excluding genes with no indication of driver potential thus lessening the false-discovery burden. Our analysis yielded 39 significant promoter hotspots at FDR<25% (Fig 3e), of which 19 in non-skin tumors and 20 in skin; there are additional 17 hotspots (5 non-skin and 12 skin) in FDR 25-50% range. Being aware that in skin there exist high mutation risk-hotspots near TSS due to UV damage and/or repair exclusion, we have, conservatively, in the skin analysis excluded TSS-proximal hotspots containing the hypermutable TTCGG motif^8^ although we do not rule out individual instances thereof could be selected. The skin WGS hotspots analysis appears reliable overall in its control for confounding mutation risk, because we did not note more inflation in the genome-wide analysis of skin compared to other cancer types (Supp Fig 8).

To rigorously assess potential effects of the promoter hotspots, in either non-skin or in skin cancers equally, we applied the PromoterAI tool^82^ to a subset of the promoter mutations at distances up to 500nt of the TSS (Fig 6a). To establish an operational definition for an impactful mutation, we used the 2.5th and the 97.5th percentile of the distribution of PromoterAI scores for the TSS-proximal mutations in passenger genes (shown as dashed lines in Fig 6a). Various *TERT* promoter mutations are ranked highly by both our mutation recurrence (selection) FDR and also by PromoterAI, serving as a positive control. All *TERT* selected promoter mutations we observed here were also reported earlier^71^ (shown in Supplementary Fig. 10). The −149 hotspot we find among the selected hotspots (at FDR<25%), although this was earlier reported as “atypical” *TERT* promoter mutation. This implied a likely passenger that occurs as part of a single, clustered-mutagenesis event, containing a driver *TERT* promoter mutation^71^, therefore our observation of it being apparently selected can be explained -- unusually for somatic mutations -- by genetic linkage. Moreover the 5’ UTR hotspots (Fig 3e) in principle have potential to perturb promoter activity therefore we also consider their TSS-activity altering potential in *TERT*. Interestingly, we identified 4 hotspots downstream of the TSS under selection (FDR<25%), two of which at +15 and +29 correspond to known kidney-specific drivers that activate *TERT* by disrupting a repressor-binding E-box motif^83^, and two others, at +1 and +13 downstream of the *TERT* TSS i.e. in the 5’ UTR we did not find reported and here identify as drivers (in LUSC+HNSC cancer type group; Supp Fig 15 and 16).

**Figure 6.**
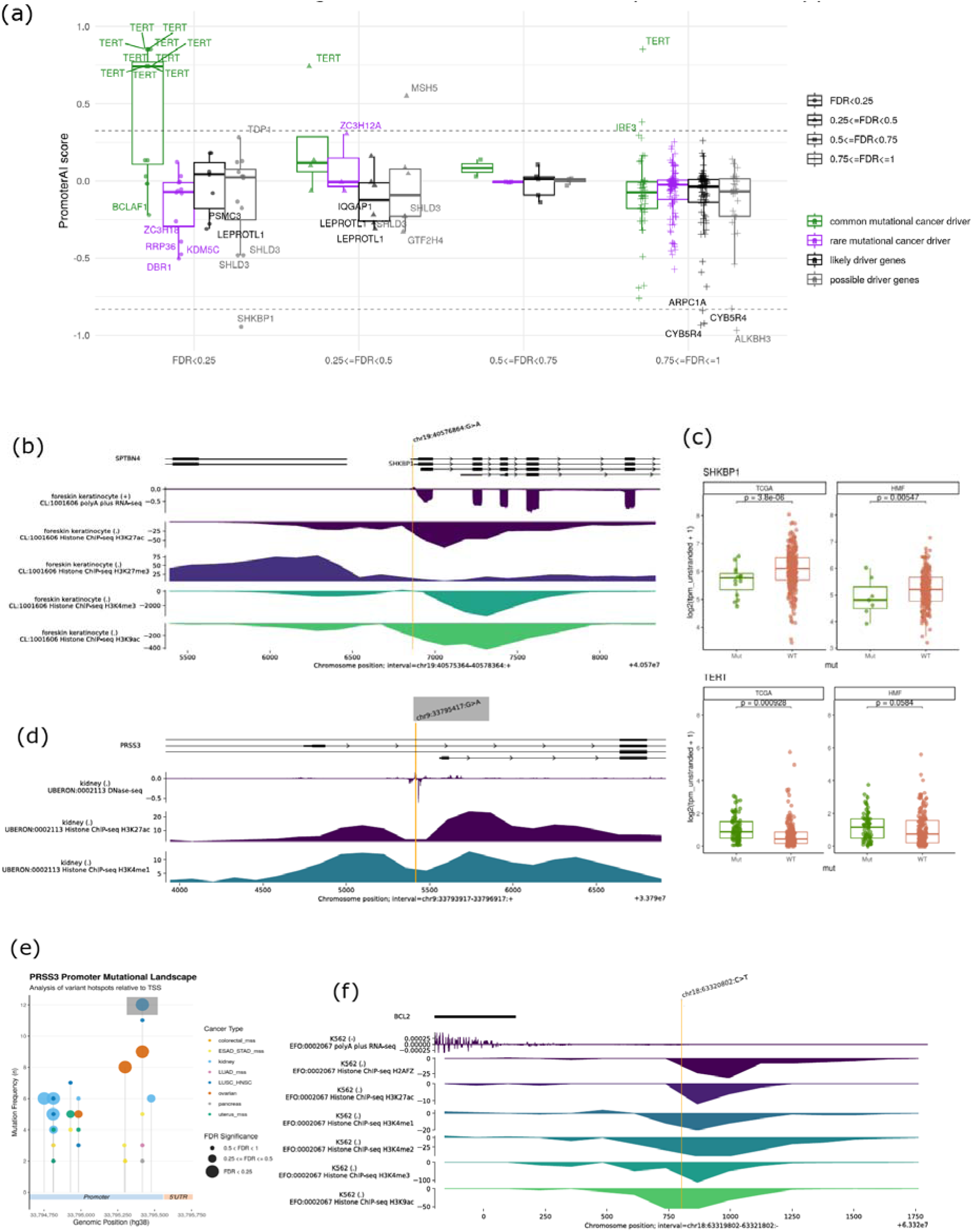
Positive selection and independent validation of non-coding mutation hotspots affecting promoters. **(a)** Distribution of PromoterAI scores across transcription start site (TSS)-proximal hotspots, here covering the [−500, 0] extended promoter region, where 0 is TSS. Dashed horizontal lines demarcate the 2.5th and 97.5th percentiles of the putatively-neutral passenger gene distribution of PromoterAI score. Statistical testing is restricted to TSS of known driver genes, and additionally of the likely cancer gene and suspected cancer gene sets. Various *TERT* driver mutations serve as positive controls (top left), with candidates stratified by FDR selection tiers, and gene names printed for genes with higher-ranking absolute PromoterAI score. **(b)** AlphaGenome epigenomic track validation for the highly recurrent *SHKBP1* promoter hotspot (chr19:40,576,864 G>A; n = 31 in melanoma cohorts), indicating functional silencing via predicted loss of active chromatin mark intensities and predicted downregulation of mRNA transcription. **(c)** Association of *SHKBP1* hotspot promoter mutations on gene expression levels across the TCGA and HMF skin melanoma cohorts, with *TERT* promoter mutations shown as a positive control. The panel shows normalized RNA-seq expression profiles (log_2(TPM + 1)) for *SHKBP1* and *TERT* pos, contrasting wild-type against mutated samples in logistic regression adjusting for 5 gene expression PCs. **(d)** AlphaGenome epigenomic track prediction for the primary TSS-proximal *PRSS3* promoter hotspot (chr9:33,795,417 G>A; n = 12 in kidney, n = 9 in ovarian cancers, plus additional cancer types), showcasing localized changes in DNase chromatin accessibility alongside sharp increases in active histone marks (H3K27ac, H3K4me1). **(e)** Local clustering of mutational landscape across the *PRSS3* promoter region spanning ~1 kb upstream of the TSS. **(f)** AlphaGenome prediction profile for the *BCL2* promoter hotspot, modeling localized alterations in chromatin activity and RNA-seq downregulation. For tracks **(b)**, **(d)**, and **(f)**, the alignment profiles aim to couple AlphaGenome tissue-specific RNA-seq and chromatin tracks to approximately matching cancer types. Additional AlphaGenome validations for promoters in genes *FGFR2*, *SHLD3*, *KDM5C*, *SMC6* and *DBR1* are shown in Supp Fig 19.

### Multiple lines of evidence for functional effects of *SHKBP1* selected promoter hotspots

There was one other promoter hotspot mutation matching confidence of *TERT* mutations: those in the *SHKBP1* gene (chr19:40,576,864:G>A, FDR=7.6%, recurrent in n=31 of the examined skin cancers). Not only does the *SHKBP1* PromoterAI effect exceed all examined TERT promoter mutations (Fig 6a), with opposite direction of effect, but also AlphaGenome predictions support that these mutations silence the *SHKBP1* mRNA levels and reduce the activating histone marks at or downstream of the promoter (Fig 6b). Next, we tested for association of *SHKBP1* promoter mutations and RNA-Seq expression levels, in the melanoma skin cancers in the TCGA and in the Hartwig Medical Foundation (HMF) cohorts. Indeed we found significant associations replicating in both cohorts, with p=3.8e-6 and 5.4e-3, respectively (Fig 6c; regression with adding gene expression PC1-PC4 as covariate). Strikingly, the effect sizes of *SHKBP1* promoter mutations on RNA-seq levels were notable, and the p-values exceed those of *TERT* promoter mutations in the same skin cancer cohorts (p=9.2e-4 and 5.8e-2 for TCGA and HMF, respectively; *TERT* in Fig 6c). *SHKBP1* gene is not a known driver gene in our lists, however we classed it among possible driver genes, because of known roles in EGFR endocytosis and degradation. In particular, the *SHKBP1* protein binds CIN85 (*SH3KBP1*) to sequester it away from the CBL-CIN85 complex with roles in protein trafficking. From the direction of the effect of the driver *SHKBP1* promoter mutations, counterintuitively, we infer *EGFR* degradation to increase, suggesting complex mechanisms of mutagenesis where reduced *EGFR* signalling would be beneficial for the melanoma. A global gene expression analysis of *SHKBP1* promoter mutant melanoma samples, focussing on genes consistently associated in the TCGA and HMF cohorts with *SHKBP1* mutations, highlights an increased immune-hot phenotype, and higher invasiveness gene expression signatures (Supp Fig 17).

### Selected hotspots in promoters of various genes including *PRSS3* (mesotrypsin) and *BCL2* display notable driver potential

Several other significant hotspots that had support in PromoterAI (albeit not quite reaching *SHKBP1* or *TERT*) were those in promoters of DNA repair genes *TDP1* and *MSH5* (at FDR<25% and 50%, respectively), the *ZC3H12A* gene encoding the Regnase-1 regulator of immune homeostasis and inflammation (FDR<50%); Fig 6a. Further more modest PromoterAI effects, approaching the boundaries of the neutral zone (defined via passenger gene mutations, see above) were predicted for various FDR<25% promoter hotspots: in *IQGAP1* scaffolding protein that binds to targets in several key cancer signalling pathways^84^, *FGFR2* oncogene and *ASXL1* tumor suppressor, *SMC6* genome integrity protein, *DBR1* splicing factor, *SHLD3* DNA repair gene and *KDM5C* chromatin modifier (Fig 6a and Supp Fig 18). For these genes, AlphaGenome validations of mutation effect on promoter activity yielded variable outcomes, highlighting how non-coding variant effect predictors are still maturing and their reliability is variable^85,86^. As well-supported examples in AlphaGenome, of the ones mentioned above, we would tentatively prioritize *FGFR2*, *SHLD3*, *KDM5C*, *SMC6* and *DBR1* (Supp Fig 19).

A remarkable example were the clustered hotspots of the promoter of the gene *PRSS3*, encoding mesotrypsin, a trypsin variety uniquely characterized by its resistance to natural serine protease inhibitors. *PRSS3* promoter had 2 hotspots in ovarian cancer and 4 in kidney cancer, significantly selected at FDR<25%, and at 25%<=FDR<50%, one uterus-MSS hotspot, one in ovarian and two in kidney, spanning roughly 1 kb upstream of the TSS (Fig 6e; there were additional non-significant but n>=5 recurrent hotspots in LUSC/HNSC group, and in gastroesophageal cancer). AlphaGenome predictions of the most recurrent and closest hotspot to the TSS (–144 bp) at <u>chr9:33,795,417:G>A</u> (n=12 in kidney, n=9 in ovarian) supports increased levels of active histone marks as well as local changes in chromatin accessibility (Fig 6d). The remaining recurrent positions within the *PRSS3* promoter at the FDR < 25% threshold were located at greater distances from the TSS (–261 bp, –748 bp, –750 bp, and –810 bp). In contrast to the TSS-proximal hotspot, AlphaGenome representations for these distal variants showed weaker predictions for active histone marks and chromatin accessibility (Supp Fig 20). The *PRSS3* promoter hotspots, because they are locally clustered, have additional confidence that they are *bona fide* selected mutations covering an oncogenic functional element. A priori, it is unlikely they are local mutagenic risk hotspots, because they are shared across three cancer types (kidney, ovary, uterus-MSS) that have quite distinct mutational signatures, as opposed to e.g. APOBEC hotspots in DNA hairpins that can be shared across cancer types. Given the known molecular roles of *PRSS3*/mesotrypsin and prior work on cancer models perturbing its activity, *PRSS3* would plausibly benefit the tumor as a pro-invasive/pro-metastatic gene^87–89^.

Finally, we identified a similarly striking cluster of non-coding mutations within the promoter region of the oncogene *BCL2*, a well-established driver in hematological malignancies. Specifically, we mapped 7 distinct recurrent mutations (with n>=3 recurrences, of that 3 mutations with n>=4) restricted to the lymphoid cohort (Supp Fig 21) mainly in the range [−800, −1,200] bp distance from TSS. Among these, a prominent hotspot is located 1,030 bp distal to the TSS (chr18 63,320,802:C:T n=6, FDR=9%). Functional evaluation via AlphaGenome supported the regulatory activity of this cluster, yielding pronounced signal intensities for H3K4me3, indicative of a highly active chromatin state at this regulatory element (Fig 6f). While the *BCL2* promoter region is an off-target activity locus of aberrant somatic hypermutation in lymphoma^19,90^, a prior report indicated causal somatic mutations ~190nt downstream of TSS that prevent binding of a repressive factor^91^. Along these lines, also our analysis, identifying TSS-upstream putative drivers, provides an example of how the locus being hypermutable does not automatically disqualify all mutations therein from being selected. A very recent study^92^ reported that somatic mutations within the broader *BCL2* promoter region (both TSS downstream i.e. in 5’ UTR but also TSS-upstream), considered collectively, associate with higher *BCL2* mRNA levels in lymphoid malignancies; here we prioritize individual hotspots in the TSS as likely driver changes.

## Discussion

Whole-genome sequencing has repeatedly shown that the noncoding genome contains abundant somatic mutations, but comparatively few noncoding driver mutations have been established with confidence^1,9^. This has created a central paradox in cancer genomics. Regulatory DNA clearly can drive cancer when altered, as exemplified by *TERT* promoter mutations^55,93,94^, tentatively also *CDC20* and *FOXA1* promoter mutations^72,73,95^, enhancer hijacking, splice-disrupting variants and mutations affecting transcription-factor binding. Moreover various germline non-coding variants consistently confer cancer risk, across multiple tumor types^96,97^. Yet large-scale somatic WGS studies have also shown that many recurrent noncoding hotspots are passengers generated by local mutational processes rather than positive selection^7,9,15^. The major unresolved problem has therefore not been whether noncoding drivers exist, but how to distinguish them from the much larger set of recurrent passengers.

Establishing the “expected” number of passenger mutations is the challenge in noncoding driver discovery, because localized mutational phenomena associated with various mutational signatures including APOBEC, UV, Signature 17 and others ^8–10,17,18,54,98^ can create striking hotspots at individual bases and thus mimic selection. MutFormer turns the activities of mutational signatures from confounders into modelled quantities. By learning mutation risk separately for individual SBS signatures from the extended DNA sequence surrounding each base, it provides a process-aware, base-pair-resolution estimate of passenger recurrence. Instead of filtering known problematic hotspots after discovery, we directly model the local sequence grammar that makes such hotspots likely to occur.

MutFormer also advances the use of sequence AI in cancer genomics. Typical genomic AI models predict variant function: effects on chromatin, transcription-factor binding, regulatory activity or gene expression^44–46,56,82^, and so they answer a different question from the one posed by driver discovery. A “functional” variant in a biochemical sense is not necessarily a selected variant; passengers can alter molecular readouts without contributing to tumor fitness. MutFormer instead models the probability that a variant would arise under neutrality. We then combine this selection-oriented AI model with functional predictors such as SpliceAI^99^, PromoterAI^82^ and AlphaGenome^76^. This separation of tasks—first asking whether recurrence exceeds mutability, then asking whether the candidate has a plausible molecular effect—aims to provide a more rigorous framework for interpreting noncoding hotspots.

The demonstration that extended sequence-dependent mutability is widespread across somatic mutational processes (MutFormer indicates ~20-mer effects for many signatures) helps explain why recurrent hotspots can arise in the absence of selection and why fixed trinucleotide or kilobase-scale background models remain vulnerable to false positives. It also aims to place various known anecdotal examples of localized mutability into a unified framework.

When applied to an expansive WGS cohort, this framework supports a conservative view of the noncoding driver landscape. Most recurrent noncoding hotspots, including many with apparently impressive recurrence, are compatible with passenger mutability^9,42,55^. However, our analysis also argues that once the passenger background risk is controlled, selected noncoding hotspots, while rare, are concentrated near known or plausible cancer genes, thus allowing to assess the prevalence of non-coding drivers.

The clearest class of such candidates in our study involves splicing. We identified selected splice-region (i.e. outside of the canonical splice site dinucleotide) or deep-intronic hotspots in several cancer genes, including *TP53*, *VHL*, *APC*, *PBRM1*, *PTPRT*, *BCL6*, *PTEN* and *TCF7L2*. Some are expected or previously recognized, validating the approach; others appear to nominate less appreciated somatic mechanisms. The deep-intronic *BCL6* hotspot in lymphoid cancers notably is predicted to generate an alternative splice donor site in a gene central to B-cell malignancy. The *PTEN* deep-intronic hotspot, supported by splice predictions and germline pathogenicity evidence, suggests a somatic route to cryptic exonization. The paired *TCF7L2* intronic hotspots in colorectal cancer point to altered splicing of an alternatively used exon in a Wnt pathway effector. These examples illustrate how important noncoding drivers may reside outside canonical promoter/enhancer annotations and outside the immediate splice dinucleotide, requiring both accurate recurrence modeling and variant-effect interpretation.

Promoter and TSS-proximal hotspots form a second major class. We recover *TERT* regulatory hotspots, including 5′ regulatory events, as a positive control for the framework. More importantly, we nominate additional promoter candidates with multiple lines of support. The *SHKBP1* promoter hotspot in melanoma shows evidence of positive selection, strong predicted promoter impact, AlphaGenome support for reduced promoter activity, and replicated association with lower *SHKBP1* expression in independent melanoma cohorts. PRSS3 shows clustered promoter hotspots across kidney, ovarian and uterus cancers, with predicted effects on active chromatin at the strongest TSS-proximal event; the recurrence of multiple nearby promoter mutations across tumor types with distinct mutational processes supports a selected regulatory element rather than a single mutational artifact. In lymphoid cancers, *BCL2* promoter-region hotspots illustrate an important principle: a locus can be hypermutable and still contain individual mutations under selection. Mutability should not automatically disqualify a candidate; instead, it should be quantitatively included in the null model.

Our study supports the view that noncoding cancer drivers have been difficult to discover because they are less frequent, more context-dependent and more dispersed than coding drivers. Even with more than 18,000 tumor WGS, many medium-recurrence hotspots remain underpowered for individual significance testing. Aggregate analyses suggest that a modest fraction of medium-recurrence promoter and other noncoding hotspots in cancer genes may be selected, implying a long tail of weak and/or tumor-type-specific regulatory drivers. Power issues will be alleviated by larger WGS cohorts, and by restricted hypothesis testing that relies on improved tumor-type-specific regulatory maps. More importantly however, we suggest the next phase of noncoding driver discovery will depend both on precise mapping of local mutational risk using improved approaches analogous to our or related methods, and also on better inferences of regulatory function of somatic variants using sequence-to-function models.

## Methods

### Patient cohorts, data scale, and study design

This study was executed via a two-phase genomic pipeline designed to first train high-resolution baseline mutation models and subsequently scale up to map positive selection across an expanded clinical cohort.

-Phase 1 (Discovery and Model Training): An initial discovery cohort comprising whole-genome sequencing (WGS) data from 12,166 tumors was assembled to extract robust *de novo* mutational signatures and train our sequence-dependent deep learning architecture (MutFormer). This training set integrated aggregated, high-yield somatic mutation calls from international repositories, including the International Cancer Genome Consortium data release (ICGC, n = 4,825, which historically encompassed the Pan-Cancer Analysis of Whole Genomes [PCAWG]^100^ joint umbrella), the Hartwig Medical Foundation^101^ (HMF, n = 4,820), the Clinical Proteomic Tumor Analysis Consortium (CPTAC, n = 1,147), a preliminary core subset of The Cancer Genome Atlas (TCGA, n = 717), the Multiple Myeloma Research Foundation CoMMpass study (MMRF-CoMMpass, n = 552), and DECIDER^102^ (n = 105).

-Phase 2 (Hotspot Extension, Cross-Consortium Harmonization, and De-deduplication): To maximize statistical power for the base-pair resolution hotspot analysis and avoid sample inflation, candidate cohorts were scaled up to include a newly consolidated, expanded release of TCGA (v43, May 2025) whole-genome architectures along with additional multi-center projects. Crucially, a strict cross-consortium harmonization and de-deduplication pipeline was applied to samples previously overlapping under the PCAWG/ICGC historical datasets. Tumors were programmatically re-assigned to their true foundational consortium-of-origin. This rigorous re-allocation shifted cross-mapped PCAWG historical specimens into the unified TCGA matrix, resulting in a finalized non-overlapping analytical cohort of 18,317 unique WGS tumors stratified across 11 distinct international projects: TCGA (n = 6,444), Hartwig Medical Foundation (HMF, n = 4,820), MUTOGRAPHS^103^ (n = 1,535), OCCAMS (n = 400), ICGC (strictly non-US international cohorts, n = 1,330), CPTAC (n = 1,147), CRC SW ^104^(n = 1,063), CoMMpass (COMMPASS, n = 906), DECIDER (n = 328), POG 570^105^ (n = 211), and OVCARE^106^ (n = 133). This unified landscape encompasses approximately 600 million single-nucleotide variants (SNVs).

### Mutational signature extraction and single-mutation assignment

*De novo* mutational signatures were extracted across the Phase 1 cohort (n = 12,166 WGS tumors) utilizing the non-negative matrix factorization (NMF) framework implemented in SigProfilerExtractor. Matrices compiling raw counts for 96 trinucleotide single base substitutions (SBS) and 83 insertion-deletion (ID) features were initially generated from raw VCF files using SigProfilerMatrixGenerator. Both matrices were combined^51^, and signature extraction was performed using default parameters across 80 NMF iterations. Based on factorization stability and reconstruction accuracy, a rank of K = 46 was selected as the largest solution maintaining an overall stability index > 0.8.

To assign specific mutational processes at individual variant resolution, *de novo* extraction profiles were decomposed into the COSMIC reference signature database (v3.4) using the decompose_fit() function from the SigProfilerAssignment package. Single-mutation probability vectors were extracted using the export_probabilities_per_mutation parameter. To ensure a highly stringent assignment and minimize false-positive process inflation, single mutations were explicitly assigned to a signature only if their posterior attribution probability exceeded 0.90. This stringent threshold yielded a core training matrix of approximately 93 million highly confident single-nucleotide variants (SNVs) distributed across 41 distinct COSMIC SBS signatures.

### Base-Pair resolution mutation rate prediction (MutFormer)

At base-pair resolution, localized mutation rate prediction was framed as a binary classification problem using a novel transformer-based neural network architecture designated as MutFormer. For each designated mutational signature, the network was trained to distinguish mutated genomic coordinates (Class 1) from non-mutated genomic control sites (Class 0). To isolate structural and motif-driven sequence patterns from baseline trinucleotide frequencies, non-mutated control coordinates were dynamically sampled from the immediate flanking windows (spanning [−1,000bp, −500bp] and [+500bp, +1,000bp] relative to the target mutation) while strictly preserving the identical 3-base trinucleotide context of the mutated site.

MutFormer processes a one-hot encoded DNA sequence of 101 nucleotides centered on the target base and outputs a single continuous scalar probability bound between 0 and 1. The model incorporates a sequential hybrid design featuring localized convolutional blocks with residual skip connections followed by a multi-headed self-attention block with rotary position embeddings.

The initial feature extraction layer consists of four sequential 1D convolutional blocks with a fixed layer width of C = 512 channels. The first convolutional block utilizes a kernel size of K = 7 with a stride of 1, dilation of 1, and symmetric zero-padding to preserve input spatial dimensions. Subsequent convolutional blocks maintain a kernel size of K = 3. Each block is structured using a Pre-Activation format consisting of a Batch Normalization layer, an Exponential Linear Unit (ELU) activation function, a 1D convolutional layer, a secondary Batch Normalization layer, a Gaussian Error Linear Unit (GELU) activation function, and a final 1×1 convolutional layer that projects features back into the residual pathway. Batch Normalization and ELU steps are omitted solely in the very first layer of the network.

To preserve precise spatial coordinate relationships across the sequence, positional information is injected into the network using Rotary Position Embedding (RoPE) layers. The transposed output fields from the convolutional stack undergo Layer Normalization and are fed into a Multi-Head Attention layer configured with 8 heads, a hidden dimension of 512, and key/value dimensions of 64. Dropout (p = 0.05) is applied directly to the computed attention matrices. Computations are accelerated by using native PyTorch FlashAttention kernels. The attention representation passes into a fully connected Feed-Forward network structured with a Gated Linear Unit (GLU) activation, Layer Normalization, dropout (p = 0.25), and a linear layer composed of 512 neurons. The final classification head consists of a sequential block featuring a Batch Normalization layer, a GELU activation, a 1×1 convolution spanning 1024 channels, a dropout layer (p = 0.1), a final GELU activation, and a single-neuron output layer with a Sigmoid activation function to generate the final prediction.

MutFormer models were trained on single NVIDIA GeForce RTX 4070 Ti or RTX 3090 GPUs, with individual signature runtimes ranging from 6 minutes to 6 days depending on total assigned mutation densities. Optimization was executed using the AdamW algorithm with a fixed batch size of 64, a base learning rate of 2e-7, and a weight decay rate of 0.0001. Models were trained for up to 100 epochs utilizing an early stopping strategy tied to a patience threshold of 15 epochs without an increase in the validation Matthews Correlation Coefficient (MCC). Data augmentation was incorporated by randomly reverse-complementing 50% of the sequences per batch to enforce genomic strand symmetry. Model hyperparameters and metrics were tracked online via the Weights & Biases platform.

Optimal hyperparameters were determined via a Bayesian hyperparameter sweep performed across signatures SBS1, SBS2, SBS10a, and SBS17b using a random sampling of 80,000 training, 20,000 validation, and 50,000 testing instances. The sweep evaluated 25 discrete architecture instances per signature across a parameter space testing channel capacity, convolutional depth, kernel combinations, transformer block layers, and dropout weights. The top two performing configurations, evaluated by the Area Under the Receiver Operating Characteristic (AUROC) on the 50,000 hold-out test set, were selected, extended across all remaining mutational signatures, and ranked globally to lock the finalized MutFormer architecture. Generalizability was evaluated by enforcing a strict cohort-based train-test split strategy where models were evaluated on entirely independent donor populations.

### Sequence determinants and structural motif interpretation

The predictive sequence context was quantified by performing *in silico mutagenesis* on the top 10 thousand most confidently predicted mutated sites in the test set per each signature. Namely, we set the importance of a position *p* in the sequence *x* to the mean change in MutFormer prediction (*MF*) for substituting all bases *b* at position *p* in *x* (*Sub*(*x, p, b*)) as:

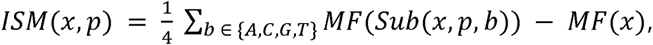

which we then average across all sequences in a given signature to get *ISM*(*p*). We treated *ISM*(*p*) ∈ *Flanks* for *p* ∈ [1,19] ∪ [81,101], and computed the effect size between *ISM*(*p*) and *Flanks* as the absolute value of Cohen’s *d*. We report positions where *d > 1* as statistically significant. Downstream de novo motif enrichment within highly scored sequences was identified using STREME via the memes R wrapper, comparing the top 100,000 scored sequences against the bottom 100,000 scored sequences. Statistical significance of motif overrepresentation was evaluated using Fisher’s exact tests, reporting both odds ratios and corresponding p-values. Discovered position weight matrices were visualized using the universalmotifs package.

### Functional genomic annotation of somatic variants

Functional annotation of somatic variants across the expanded ~600M SNV matrix was carried out using the Ensembl Variant Effect Predictor (VEP) tool. Single nucleotide variants were partitioned into discrete functional genomic categories based on their regional mapping coordinates:

- Coding: Variants explicitly annotated as *missense_variant*, *synonymous_variant*, or *stop_gained*.
- 5’ UTR / 3’ UTR: Variants matching *5_prime_UTR* and *3_prime_UTR* terms, respectively.
- Canonical Splicing Sites: Variants altering standard splice boundary domains, including *splice_donor_variant*, *splice_donor_region_variant*, *splice_acceptor_variant*, and *splice_polypyrimidine_tract_variant*.
- Cryptic Intronic Splice Sites: To capture non-canonical splicing disruptions occurring deeper within intronic regions, deep learning-based splicing evaluations were performed using SpliceAI. Variants mapping to intronic regions that exhibited a maximum delta score (representing splice donor/acceptor gain or loss) greater than a stringent threshold of SpliceAI > 0.10 were programmatically rescued and annotated as functional splicing-disrupting mutations.
- Promoters: Non-coding variants localized near transcription start sites (TSS) were partitioned into Promoter 200 (distance from TSS <200bp) and Promoter 1500 (200bp < distance from TSS < 1,500bp) based on *upstream_gene_variant* classifications.
- Enhancers: Variants falling within coordinates compiled in the validated catalog of transcriptionally active human enhancers published by Lidschreiber et al^74^.
- lncRNA: Non-coding variants overlapping long non-coding transcripts carrying the *lincRNA* biotype annotation.

### Somatic hotspot database assembly, mutational signature deconvolution, and artifact filtering

To establish a robust database of recurrent somatic mutation hotspots at single-base resolution across the expanded cohort of 18,317 whole-genome-sequenced (WGS) tumors, we developed a multi-layered filtering and analytical pipeline designed to eliminate localized variant-calling artifacts, technical noise, and mapping vulnerabilities.

Initial candidate loci were strictly restricted to genomic coordinates displaying a cumulative cross-cohort mutation count of *m* ≥ 2 within a given cancer type histology, requiring validation across at least two independent international sequencing centers to minimize center-specific biases. To minimize variant-calling false positives arising from non-unique mappings or sequence homology, the analytical space was strictly constrained to high-confidence regions defined by the Panmask framework (v2, pm151a.easy). Loci were excluded if they overlapped assembly gaps, non-chromosomal contigs, or genomic regions prone to short-read misalignment, defined as positions where 151-mers occur ≥ *n* times across the human pangenome reference collection (where *n* represents the total number of genomes evaluated).

To account for regional genomic variations in sequencing accessibility, positional mapability was further controlled; genomic positions displaying high mapability vulnerability—defined as a uniqueness score ≤ 80 across both the UMAP 24-bp and 36-bp tracks—were filtered out. Furthermore, variants overlapping known cross-assembly mapping artifacts compiled in both the GRCh37 (hg19) and GRCh38 novel Candidate Unmapped Positions (CUPs) registries, as well as ultra-high-signal artifacts captured within the ENCODE blacklist or panel-of-normals (PON) exclusionary regions, were systematically removed.

In the second phase of the pipeline, patient-specific mutational signature exposure profiles were characterized to identify and exclude sample-level technical noise. For each of the major tumor histologies, a 96-trinucleotide mutation count matrix (*X*) was constructed as an input catalog. We utilized the MuSiCal^107^ (Mutational Signature Calculator) framework to perform a constrained mathematical optimization and refit the observed tumor catalogs against the standardized COSMIC v3.2 reference spectrum for single-base substitutions in WGS data (*W*). The optimization was parameterized via a naive thresholding approach:

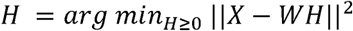

with the threshold set to 0 (thresh=0) to obtain the full compositional matrix of exposure coefficients (*H*). The resulting exposure matrices were normalized to determine the relative contribution of each signature per individual tumor.

To prevent systemic artifact inflation from confounding downstream statistical modeling, any tumor displaying an artifact or sequencing-specific signature exposure profile exceeding 10% of its global mutational burden (specifically across signatures SBS27, SBS43, SBS45–SBS56, SBS57–SBS60, and SBS95) was excluded from the database. This multi-layered filtering pipeline yielded a clean, core analytical hotspot matrix evaluated across major tumor histologies for subsequent zero-truncated Poisson regression modeling.

### Statistical hotspot modeling via target count transformation

Because genomic positions with fewer than two mutations (*m* < 2) were structurally excluded from the hotspot database due to our rigorous recurrence criteria, the raw response variable was truncated below 2. To model this left-truncated distribution within the Generalized Linear Mixed Models via Template Model Builder (glmmTMB^108^) R package, a programmatic count transformation was applied to the observed data. The raw mutation count (*m*) at each position was shifted downwards by subtracting a unit value (*n* = *m* − 1), mapping the original domain *m* ∈ {2, 3, 4, …} onto a transformed domain *n* ∈ {1, 2, 3, …}.

This transformation allowed the shifted mutation count (*n*) to be modeled via a standard Zero-Truncated Poisson (ZTP) regression framework:

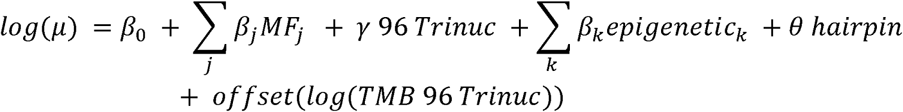

where *µ* represents the expected value of the transformed count *n*. *MF_j_* represents the continuous MutFormer signature probability, and 96*Trinuc* captures the localized 96-trinucleotide context, collapsed via proportional pooling (fct_lump_prop) to group rare contexts representing <1\% or <5\% of the data within low-density cohorts.

Confounding regional genomic factors were controlled by incorporating a comprehensive covariate matrix of localized epigenetic marks (*epigenetic_k_*), including replication timing, DNase hypersensitivity, histone modifications (H3K27ac, H3K36me3, H3K4me3, H3K9me3), DNA methylation rates, nucleosome occupancy (MNase-seq), and CTCF/RAD21 transcription factor binding intensities. Continuous track values for DNase and histone marks were winsorized at a maximum threshold value of 4 to prevent model distortion by extreme local outliers. For APOBEC-enriched tumor types (e.g., bladder and breast cohorts), an indicator variable tracking predicted fold-back hairpin structures (*hairpin*) along with its explicit interaction terms with signatures SBS2 and SBS13 was integrated into the model formula. Structural variation in specimen-specific background mutation densities was adjusted by applying a multi-variable offset calculating the log-transformed cumulative background mutation burden (*log*(*TMB* 96 *Trinuc*)) across all tumor samples harboring mutations at that specific 96-trinucleotide context.

### Iterative residual analysis and outlier removal

To ensure that baseline passenger rate models were not systematically biased by highly recurrent driver hotspots undergoing intense positive selection, an iterative residual desensitization pipeline was implemented during model optimization. The initial training dataset was structurally depleted of known cancer drivers by removing coordinates mapping to curated driver genes in the MutPanning and Cancer Gene Census (CGC) databases.

An initial ZTP model was fitted on this clean training set, and absolute residuals (|*n* − *m*|) were calculated for all coordinates. To smooth out extreme localized overdispersion without erasing volatile recurrent regions, rows displaying the top 50% largest absolute residuals were iteratively pruned. Pruning was controlled by a logical check to ensure that the deletion did not eliminate the final remaining representative row of a multi-row hotspot cluster. Models were re-fitted dynamically using a protective execution function to catch non-convergence or singular matrix errors. This cleanup cycle was repeated until all multi-row hotspots were reduced to a stable configuration, followed by a final re-fitting under the strict ZTP parametrization to yield the unbiased baseline passenger expectations.

### Positive selection testing and non-canonical hotspot discovery

To uncover non-canonical genomic loci under active positive evolutionary selection, expected value predictions (*µ*) derived from the final passenger glmmTMB models were re-shifted back to the original biological scale (*E* = *µ* + 1) and compared directly against raw observed cross-cohort mutation counts (O, where O = n + 1). The statistical significance of positive deviations (O>E) was formally evaluated using a one-degree-of-freedom Chi-square (*X^2^*) goodness-of-fit test:

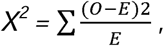

Positions where the observed mutation count fell below the baseline model expectations (O>E) were assigned a p-value of NA. Multiple testing corrections were executed across the genome utilizing the Benjamini-Hochberg False Discovery Rate (FDR) procedure. To ensure robust statistical power and control for Type I error propagation in low-density contexts, multiple testing adjustments were restricted to hotspots displaying an observed count of O>3 or O>4 in extremely mutated cancer types such as colorectal MSI and colorectal+uterus POLE.

The accuracy of the combined prediction framework was validated by compiling the Area Under the Receiver Operating Characteristic curve (AUC-ROC) via the pROC package, contrasting the relative prediction profiles of validated coding driver genes against baseline passenger backgrounds.

### Multi-layered orthogonal validation of selected hotspots

To independently validate the functional impact and oncogenic potential of the discovered non-canonical hotspots under positive selection, we implemented a multi-layered benchmarking pipeline utilizing computational variant-effect predictors, structural modeling frameworks, and curated clinical variant registries:

- Coding variant validation via AlphaMissense: The functional impact of recurrent somatic missense alterations identified within positive-selected hotspots was systematically cross-referenced with AlphaMissense pathogenicity scores.
- Non-canonical splicing verification via SpliceAI and ClinVar: Beyond the initial filtering of canonical splice coordinates, candidate hotspots localized within intronic boundaries or cryptic splicing junctions were rigorously benchmarked. We evaluated the specific splice site creation or disruption dynamics using SpliceAI delta scores. To provide direct clinical orthogonality, the predicted splicing-disrupting hotspots were queried against the ClinVar database. Hotspots overlapping known pathogenic or likely pathogenic variants associated with aberrant splicing mechanisms were designated as high-confidence splicing drivers.
- Non-Coding promoter disruption modeling via PromoterAI. The regulatory disruption potential of recurrent alterations localized within transcription start site windows (Promoter 200 and Promoter 1500) was evaluated using PromoterAI.
- Non-Coding regulatory validation via AlphaGenome plots. To model the comprehensive sequence-to-phenotype alterations induced by non-canonical variants outside of standard coding regions, we utilized AlphaGenome, a unified 1-Mb context window deep-learning architecture capable of multi-track operational predictions at base-pair resolution. Candidate positive-selected hotspots were comprehensively evaluated across specific genomic modalities:

- *Splicing Sites:* AlphaGenome sequence mutations were used to predict structural alterations in splice site usage, mapping precise deviations in single-nucleotide splice junction coordinates and splice strength.
- *Promoters:* For variants within Promoter 200 and Promoter 1500 windows, we quantified AlphaGenome-predicted shifts in transcription initiation, localized chromatin accessibility, and transcription factor (TF) binding profiles at the target core promoters.
- *5’ UTRs:* Functional disruption of untranslated regions was evaluated by tracking model-predicted downstream changes in localized gene expression tracks and post-transcriptional regulatory motifs.

## Code and Data Availability

In this study, published tumor datasets were reanalyzed. We obtained the WGS somatic mutation calls for the Hartwig Medical Foundation study [https://www.hartwigmedicalfoundation.nl/en/] under request DR-260 (Hartwig data is available under restricted access via https://www.hartwigmedicalfoundation.nl/en/data/data-access-request/), the PCAWG study re-processed by the HMF pipeline at the International Cancer Genome Consortium (ICGC) data portal [https://dcc.icgc.org/releases/PCAWG/Hartwig], the Personal Oncogenomics (POG) project from BC Cancer [https://www.bcgsc.ca/downloads/POG570/], the DECIDER data from within a collaborative project [https://www.deciderproject.eu/] also available under restricted access via EGA accession number EGAS00001006775. We downloaded bam files for CPTAC-3 project (available under restricted access via dbGaP accession phs001287.v17.p6 [https://www.ncbi.nlm.nih.gov/projects/gap/cgi-bin/study.cgi?study_id=phs001287.v17.p6]), and MMRF-COMMPASS project (available under restricted access via dbGaP accession phs000748.v7.p4 [https://www.ncbi.nlm.nih.gov/projects/gap/cgi-bin/study.cgi?study_id=phs000748.v7.p4]) from the NCI Genomic Data Commons data repository [https://portal.gdc.cancer.gov/]. Mutographs project is at the European Genome-Phenome archive EGAD00001006732 [https://ega-archive.org/datasets/EGAD00001006732]. We downloaded the curated WGS somatic mutation calls from Swedish colorectal patients from Nunes et al. study [accession number PRJEB61514] and from the OVCARE from Wang, Y. K. et al study. The relevant data generated in this study are provided in the Supplementary Tables or the Source Data file. The Source Data and Code to generate main Figures of this paper will be provided on GitHub upon manuscript acceptance. Code related to training MutFormer will be provided on GitHub upon manuscript acceptance.

## Acknowledgements

I.G.F was supported by Juan de la Cierva postdoctoral contract (JC2021-047350-I) from the Spanish government and Horizon2020 RIA project “DECIDER” (965193). M.V. was supported by a fellowship from the ”la Caixa” Foundation (ID 100010434, with fellowship code B006197). Group of F.S. was supported by an ERC Consolidator Grant “STRUCTOMATIC” (101088342), Horizon2020 RIA project “DECIDER” (965193), Horizon Europe project “LUCIA” (101096473), Spanish government project “REPAIRSCAPE”, CaixaResearch project “POTENT-IMMUNO” (HR22-00402), a Novo Nordisk Fonden “Start Package” grant, the Danish Cancer Society grant “AI-DRIVERS” and a DFF Project2 (5243-00072B), the SGR funding of the Catalan government, and the Severo Ochoa Centers of Excellence award to the IRB Barcelona. LLM-assisted tools were used for language editing of manuscript text.

## Author Contributions

IGF curated, processed, quality-controlled and analyzed mutation data; co-designed the analyses; contributed to the motif enrichment analysis; developed the background mutagenesis statistical framework to test for hotspot selection; collected and analyzed supporting evidence for hotspots via variant effect predictors; visualized, integrated and interpreted various data resulting from diverse analyses; drafted the manuscript. MV conceptualized and developed the MutFormer framework and trained MutFormer models; contributed to motif analysis and visualized and interpreted corresponding data; contributed to variant effect interpretation; and contributed to drafting the manuscript. DN conceptualized, developed and tested neural net architectures for modelling mutation risk; and provided scientific computing expertise. FS conceived the study; co-designed the analyses; interpreted the data, drafted the manuscript, and supervised the study.

