## Supplementary Figures 1-21 for "Transformer models of mutation risk at base-pair resolution identify non-coding hotspot cancer driver mutations"

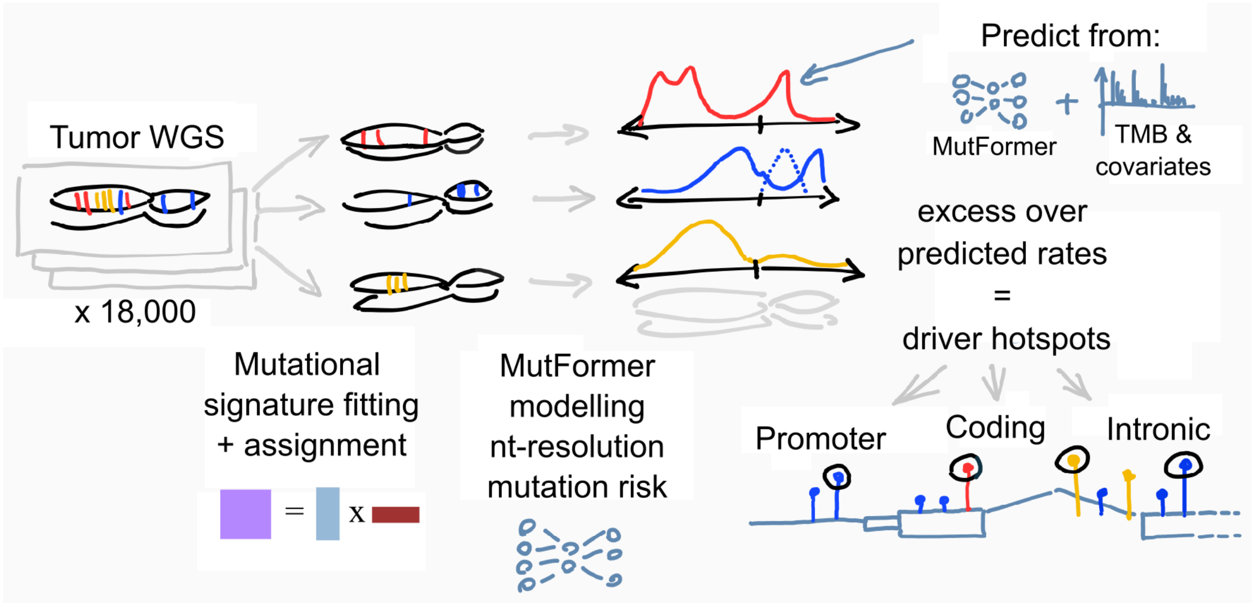


**Supplementary Figure 1. Schematic workflow of the MutFormer framework for genome-wide driver hotspot discovery.** Somatic mutation profiles from ~18,000 pan-cancer whole-genome sequencing (WGS) tumors are initially decomposed via mutational signature fitting and assignment, which factorizes the raw mutation count matrix into tumor-specific exposure vectors and distinct signature profiles. These extracted signatures are integrated with local sequence context into MutFormer, a deep-learning transformer architecture designed to model background mutation rates at single-nucleotide resolution. The model generates signature-specific expected background mutation risk tracks, represented by the non-uniform density curves. Observed local mutation frequencies are then statistically evaluated against these predicted neutral background rates. Finally, these statistically significant hotspots are functionally annotated and mapped across the genome to pinpoint driver elements within promoters, protein-coding exons, and intronic regions.

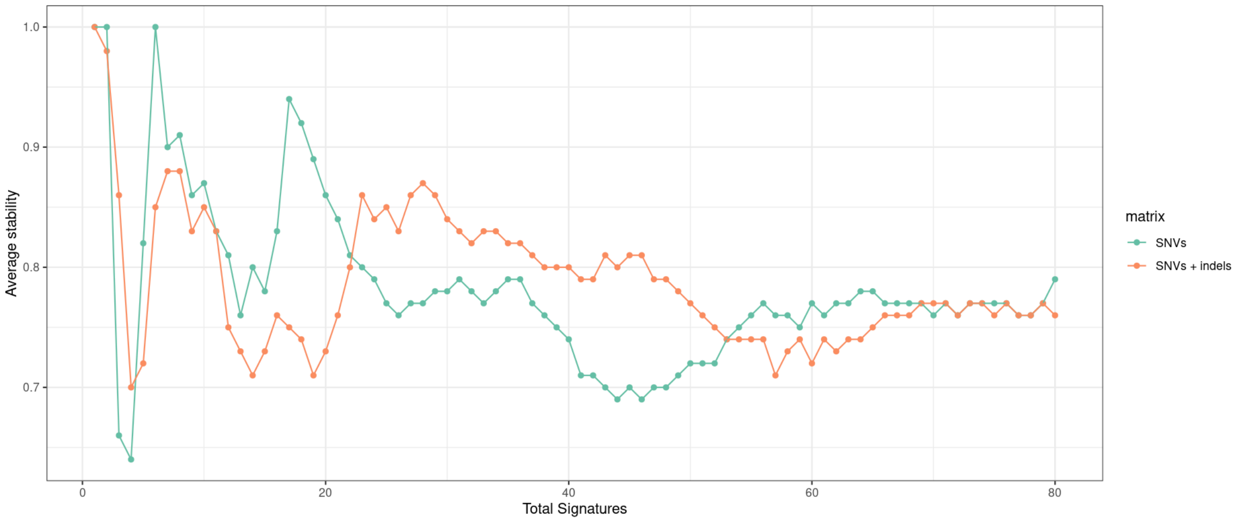


**Supplementary Figure 2. Numerical stability benchmarking of mutational signature extraction.** Lineplot evaluating the mean stability metrics of Non-negative Matrix Factorization (NMF) de novo signature extractions across varying signature extraction counts (K = 1-80 solutions). Stability performance is directly compared between NMF models trained using the standard SBS96 mutation count matrix alone (green dots) versus models utilizing a concatenated matrix combining both SBS96 and ID83 somatic indel counts (orange dots). The consistently higher stability scores achieved across K=23-50 solutions support the selection of the concatenated SBS96+ID83 matrix framework to robustly isolate independent mutagenic processes.

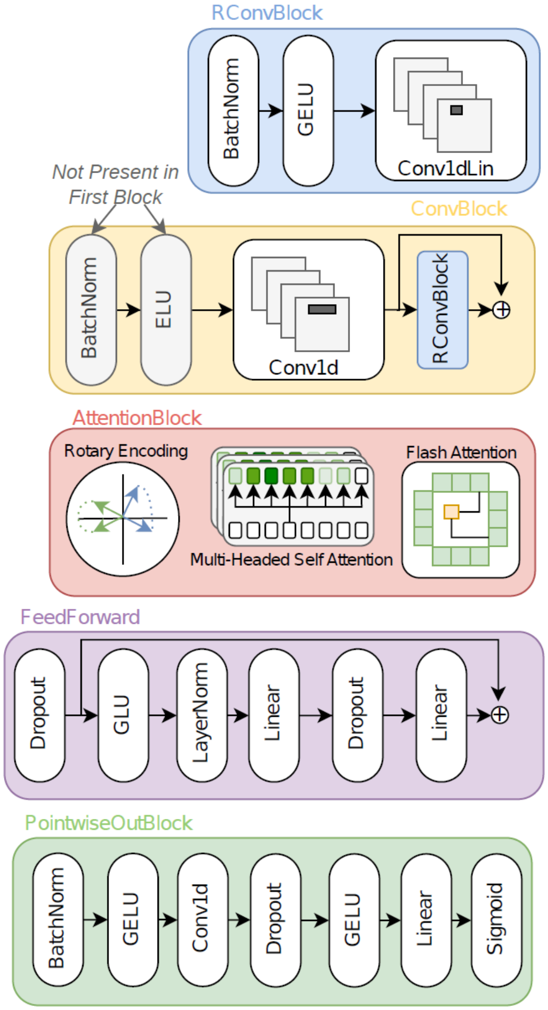


**Supplementary Figure 3. Details of the convolutional–transformer hybrid neural network architecture in MutFormer.** For each candidate site, the model takes as input a one-hot encoded DNA sequence spanning 50 bp upstream and 50 bp downstream of the central nucleotide. A series of convolutional blocks with skip connections learns local sequence motifs, followed by a multi-headed self-attention block to capture interactions between motifs across the sequence. The resulting representations are passed through a pointwise output head to produce a mutation risk score.

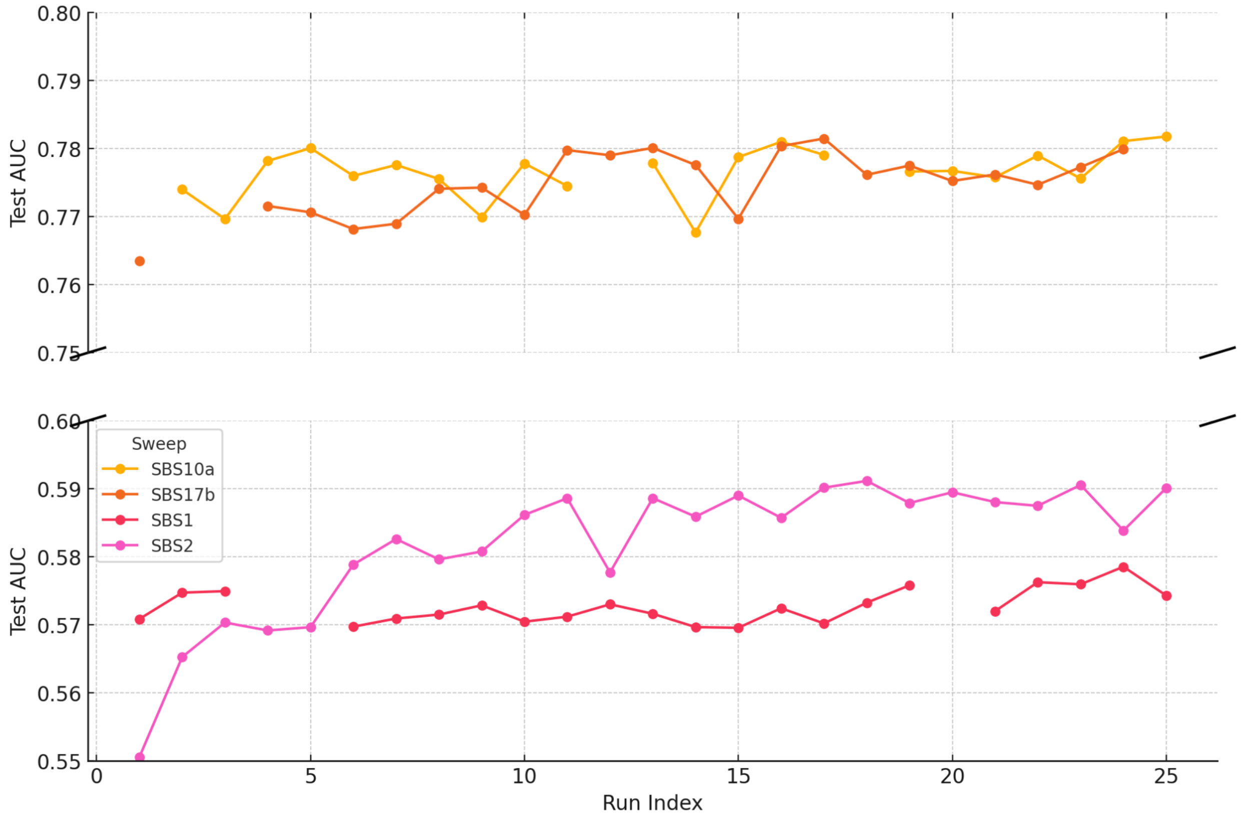


**Supp Figure 4. Hyperparameter tuning of MutFormer for selected signatures.** MutFormer performance on test data measured in AUC (y axis; higher is better) over 25 different hyperparameter configurations (x axis). Bayesian hyperparameter search or sweep was performed independently for selected signatures (SBS10a, SBS17b, SBS1 and SBS2) exploring different parameter sets. Some configurations did not converge, or resulted in GPU errors, corresponding to the missing data points.

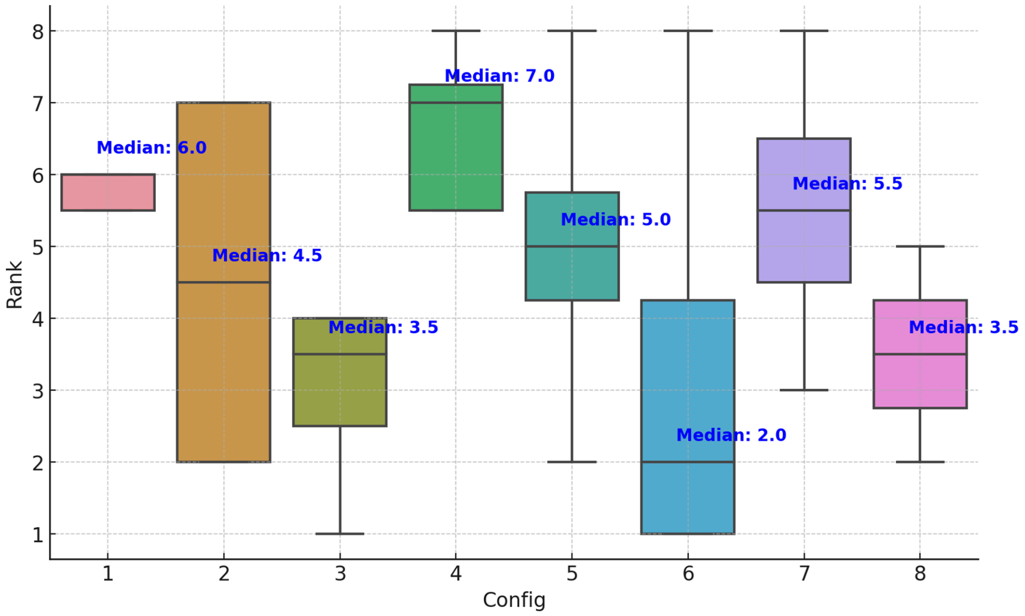


**Supplementary Figure 5. Ranking of top-performing metaparameter sets.** The top 2 parameters sets were taken by signature (SBS10a, SBS17b, SBS2, SBS1) based on Test AUC, yielding 8 configurations (x axis). The boxplot shows the rankings (y axis) of neural networks based on these 8 parameter sets across the 4 signatures, ranked by Test AUC; lower is better. The parameter set that led to the best performing model (lower rank) across the signatures was selected as final (Config 6). Boxplot colours are ornamental.

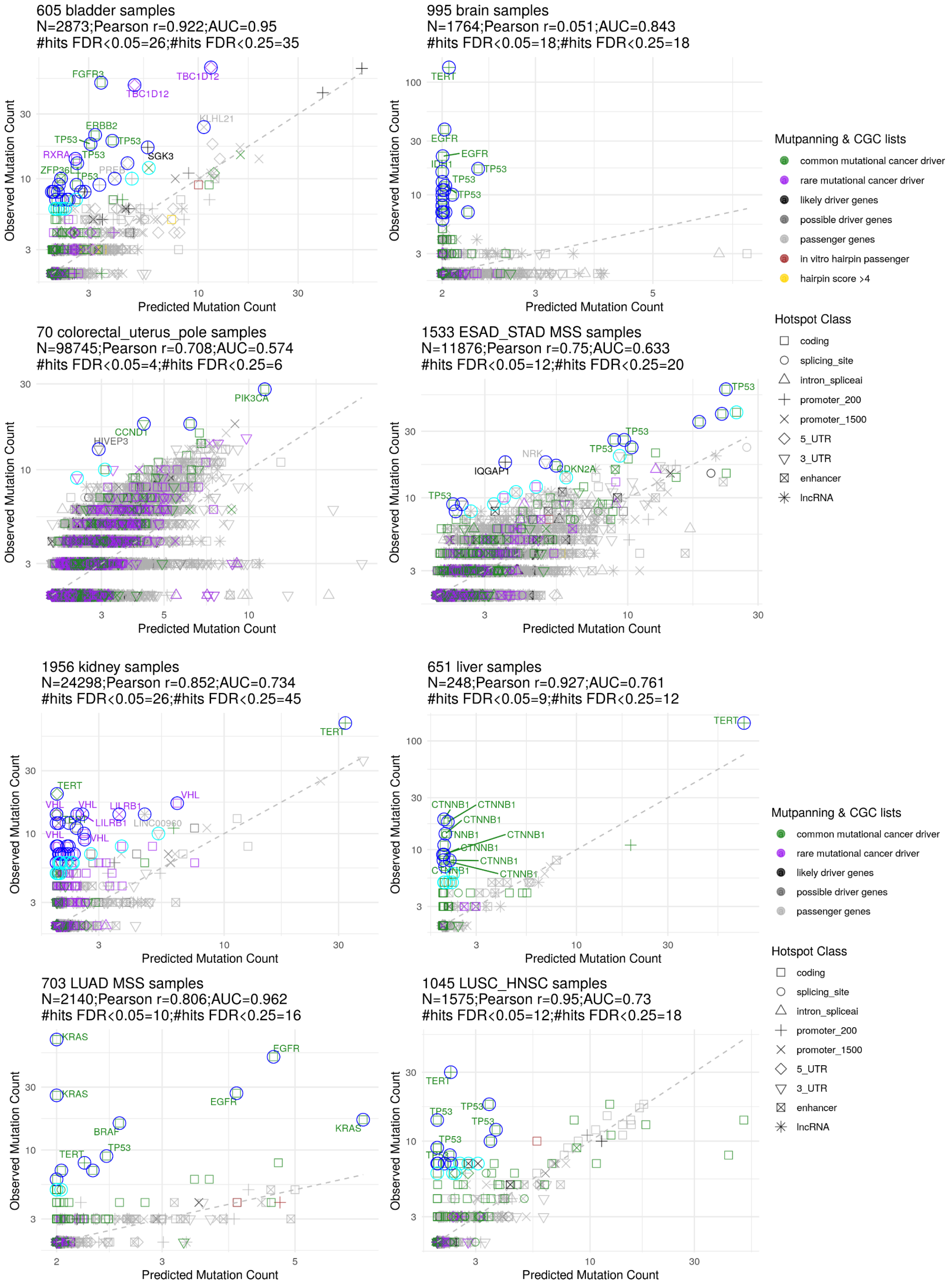
605 bladder samples
N=2873;Pearson r=0.922;AUC=0.95
#hits FDR<0.05=26;#hits FDR<0.25=35
FGFR3
TBC1D12
TBC1D12
KLHL21
ERBB2
TP53
TP53
TP53
SGK3
RXRA
ZFP36L1
TP53
Observed Mutation Count
Predicted Mutation Count
995 brain samples
N=1764;Pearson r=0.051;AUC=0.843
#hits FDR<0.05=18;#hits FDR<0.25=18
TERT
EGFR
EGFR
IDH1
TP53
TP53
TP53
Observed Mutation Count
Predicted Mutation Count
Mutpanning & CGC lists
common mutational cancer driver
rare mutational cancer driver
likely driver genes
possible driver genes
passenger genes
in vitro hairpin passenger
hairpin score >4
Hotspot Class
coding
splicing_site
intron_spliceai
promoter_200
promoter_1500
5_UTR
3_UTR
enhancer
lncRNA
70 colorectal_uterus_pole samples
N=98745;Pearson r=0.708;AUC=0.574
#hits FDR<0.05=4;#hits FDR<0.25=6
PIK3CA
CCND1
HIVEP3
Observed Mutation Count
Predicted Mutation Count
1533 ESAD_STAD MSS samples
N=11876;Pearson r=0.75;AUC=0.633
#hits FDR<0.05=12;#hits FDR<0.25=20
TP53
TP53
TP53
NRK
IQGAP1
CDKN2A
TP53
Observed Mutation Count
Predicted Mutation Count
1956 kidney samples
N=24298;Pearson r=0.852;AUC=0.734
#hits FDR<0.05=26;#hits FDR<0.25=45
TERT
TERT
VHL
VHL
VHL
LILRB1
LILRB1
LINC00960
VHL
VHL
Observed Mutation Count
Predicted Mutation Count
651 liver samples
N=248;Pearson r=0.927;AUC=0.761
#hits FDR<0.05=9;#hits FDR<0.25=12
TERT
CTNNB1
CTNNB1
CTNNB1
CTNNB1
CTNNB1
CTNNB1
CTNNB1
CTNNB1
CTNNB1
Observed Mutation Count
Predicted Mutation Count
Mutpanning & CGC lists
common mutational cancer driver
rare mutational cancer driver
likely driver genes
possible driver genes
passenger genes
Hotspot Class
coding
splicing_site
intron_spliceai
promoter_200
promoter_1500
5_UTR
3_UTR
enhancer
lncRNA
703 LUAD MSS samples
N=2140;Pearson r=0.806;AUC=0.962
#hits FDR<0.05=10;#hits FDR<0.25=16
KRAS
EGFR
KRAS
EGFR
KRAS
BRAF
TERT
TP53
Observed Mutation Count
Predicted Mutation Count
1045 LUSC_HNSC samples
N=1575;Pearson r=0.95;AUC=0.73
#hits FDR<0.05=12;#hits FDR<0.25=18
TERT
TP53
TP53
TP53
TP53
Observed Mutation Count
Predicted Mutation Count

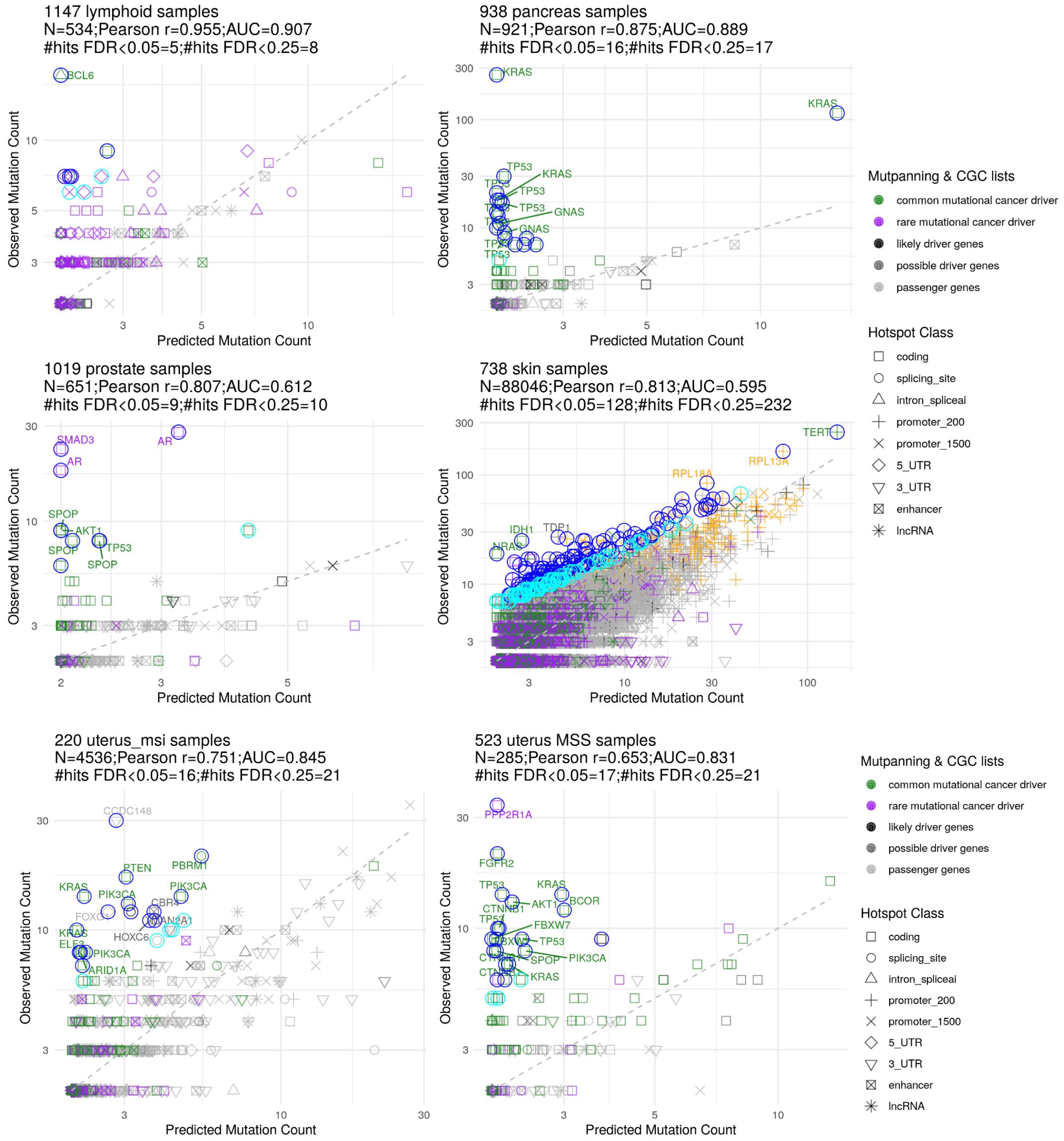


**Supplementary Figure 6.** Scatterplot of the observed mutation counts and the estimated mutation counts for **a)** bladder, **b)** brain, **c)** colorectal and uterine POLE**, d)** esophagous and stomach, **e)** kidney, **f)** liver, **g)** lung adenocarcinoma, **h)** lung squamous and head-neck squamous **i)** lymphoid**, j)** pancreas, **k)** prostate, **l)** skin, **m)** uterine MSI and **n)** uterine MSS cancer.

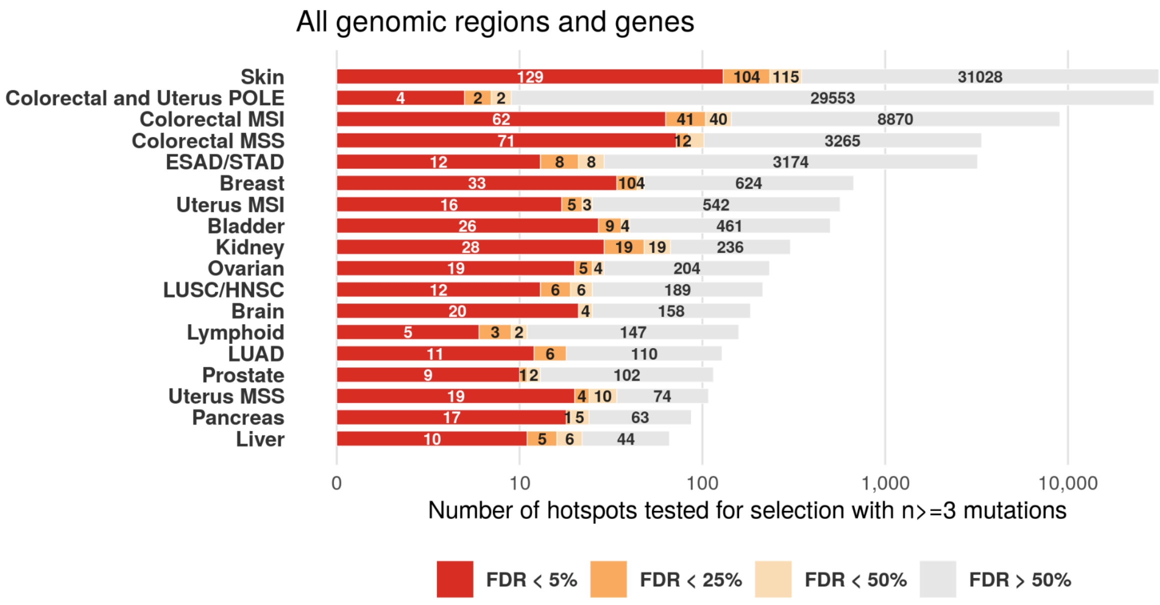
All genomic regions and genes
Skin
Colorectal and Uterus POLE
Colorectal MSI
Colorectal MSS
ESAD/STAD
Breast
Uterus MSI
Bladder
Kidney
Ovarian
LUSC/HNSC
Brain
Lymphoid
LUAD
Prostate
Uterus MSS
Pancreas
Liver
0
10
100
1,000
10,000
Number of hotspots tested for selection with n>=3 mutations
FDR < 5%
FDR < 25%
FDR < 50%
FDR > 50%

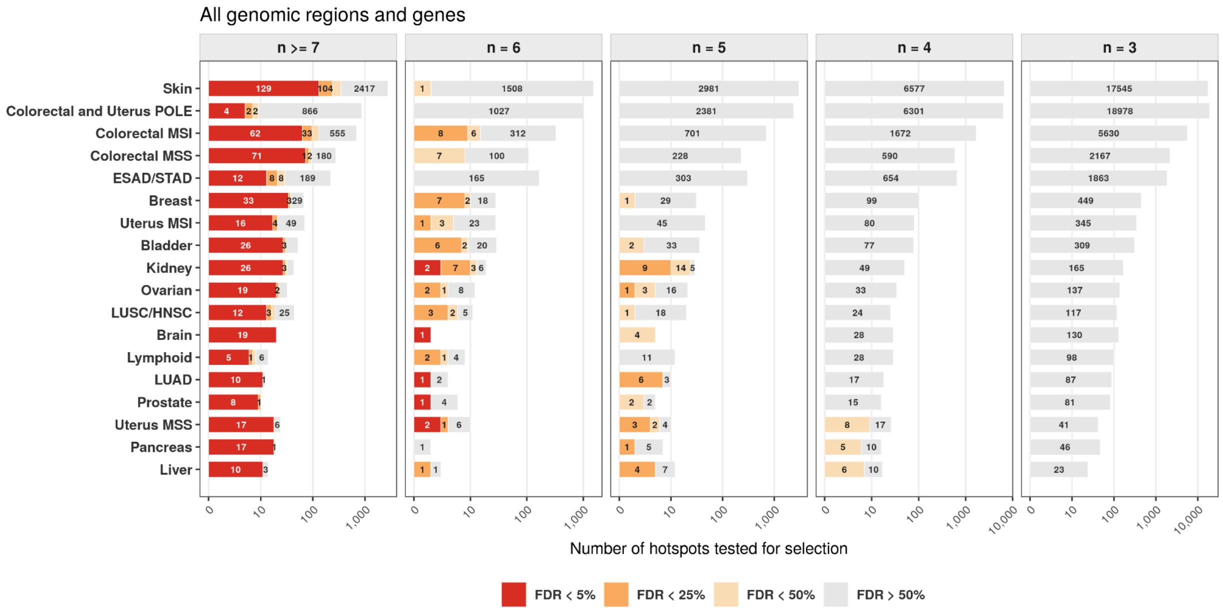
All genomic regions and genes
n >= 7
n = 6
n = 5
n = 4
n = 3
Skin
Colorectal and Uterus POLE
Colorectal MSI
Colorectal MSS
ESAD/STAD
Breast
Uterus MSI
Bladder
Kidney
Ovarian
LUSC/HNSC
Brain
Lymphoid
LUAD
Prostate
Uterus MSS
Pancreas
Liver
Number of hotspots tested for selection
FDR < 5%
FDR < 25%
FDR < 50%
FDR > 50%

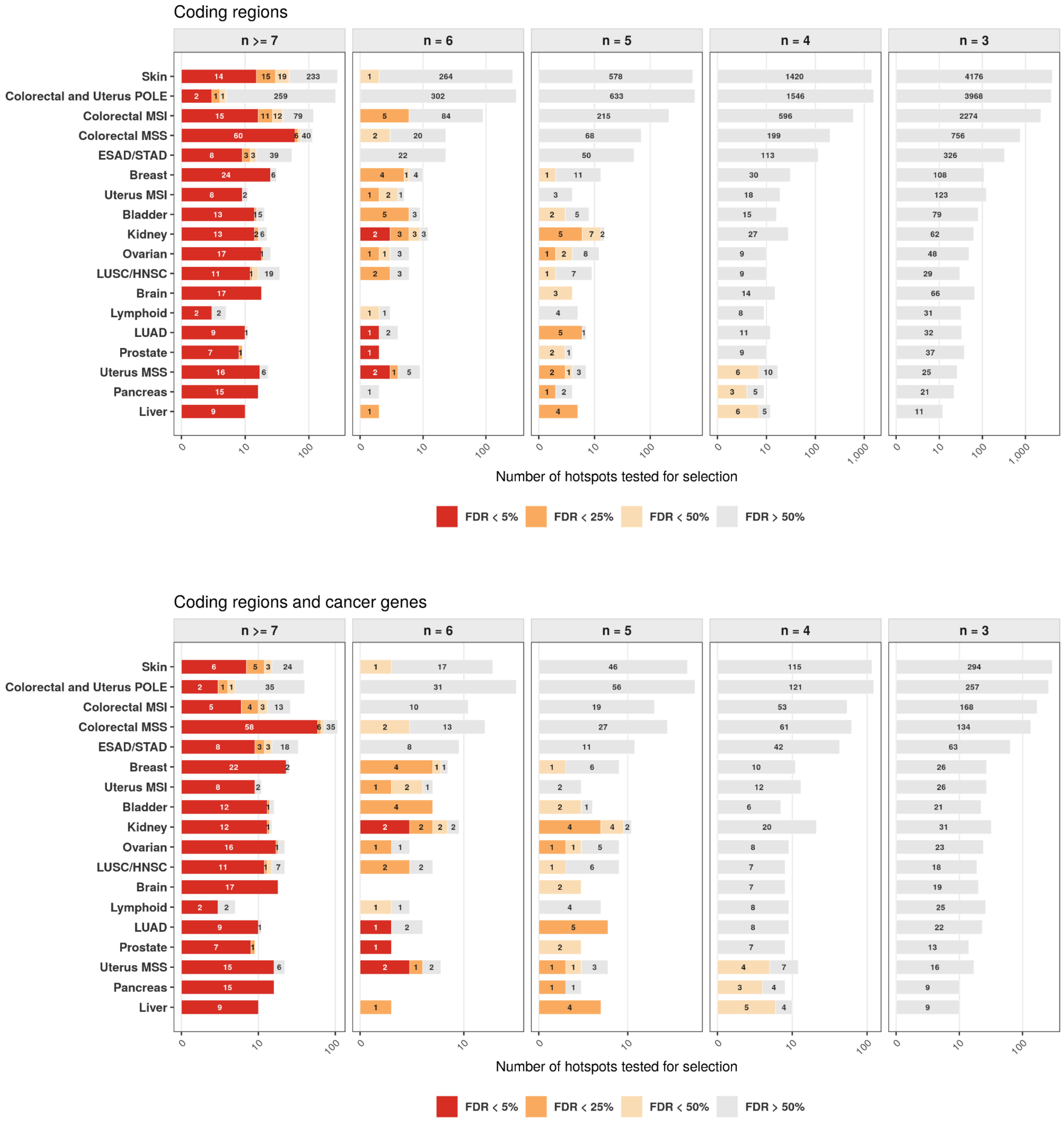


**Supplementary Figure 7.** Number of hotspots tested for selection with n>=3 mutations for each cancer type accounted by False Discovery Rate (FDR) in **a)** all genomic regions and genes; **b)** all genomic regions and genes splitted by n=3, 4, 5, 6 and >=7 mutations; **c)** coding regions splitted by n=3, 4, 5, 6 and >=7 mutations; **d)** coding regions in cancer genes splitted by n=3, 4, 5, 6 and >=7 mutations.

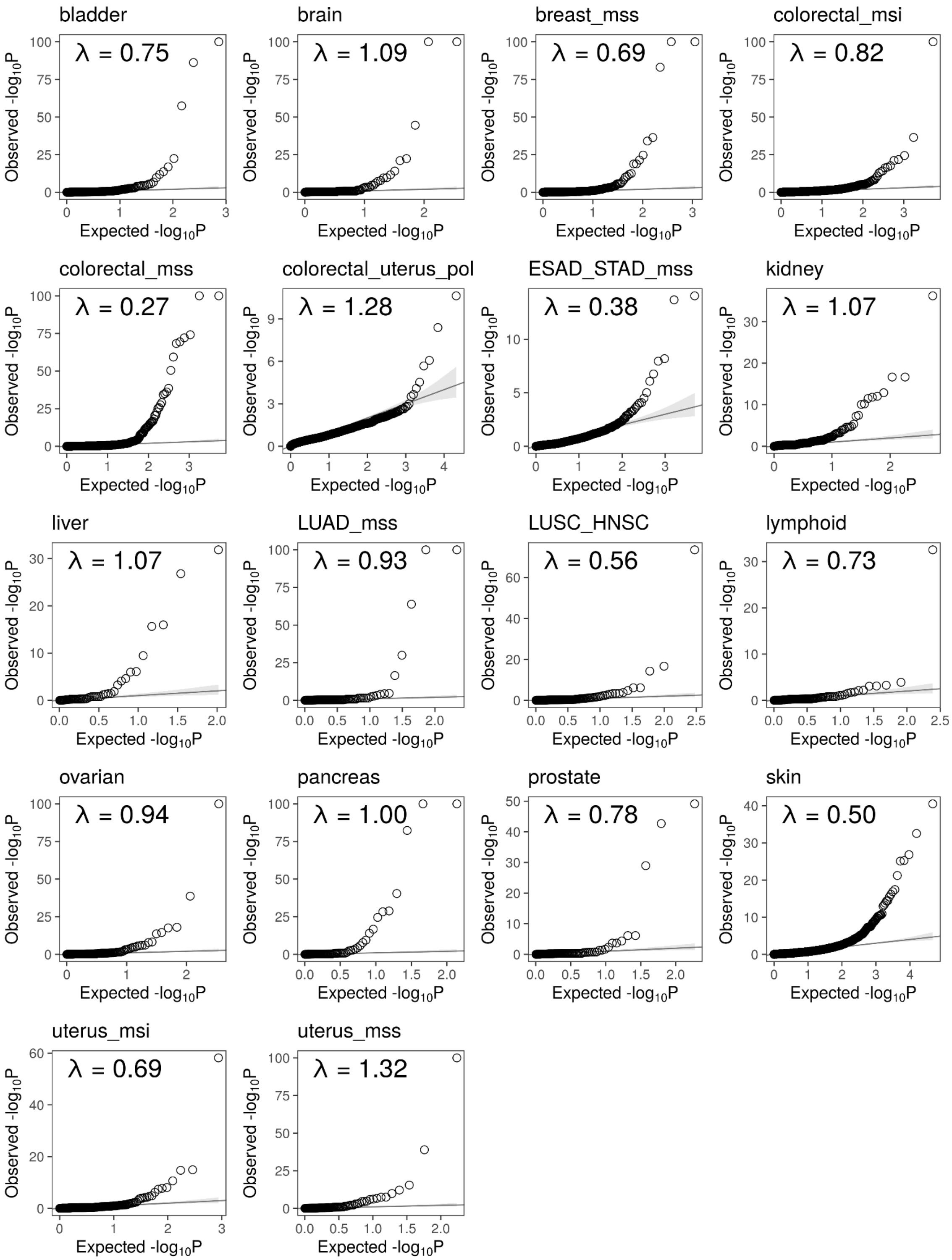


**Supplementary Figure 8. Statistical calibration of positive selection tests for genomic hotspots.** Quantile-quantile (Q-Q) plots of observed versus expected -log10(p-values) stratified by cancer type. λ denotes the genomic inflation factor for each statistical model.

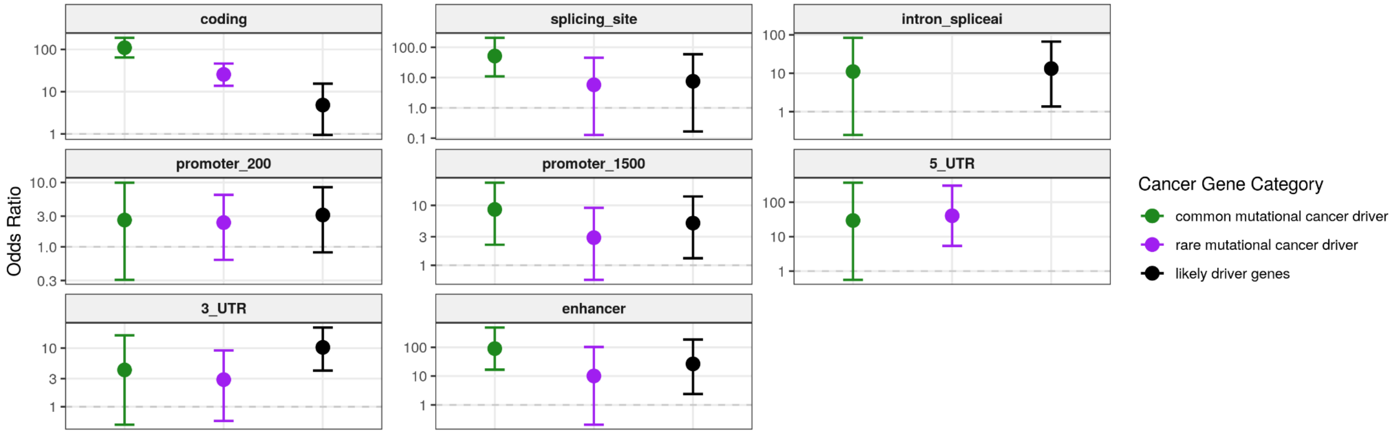


**Supplementary Figure 9. Statistical enrichment analyses of functional hotspots in cancer-associated genes.** Odds ratio estimates quantifying hotspot enrichment in common drivers, rare drivers, and likely driver genes compared to background genes. Error bars represent the 95% confidence intervals.

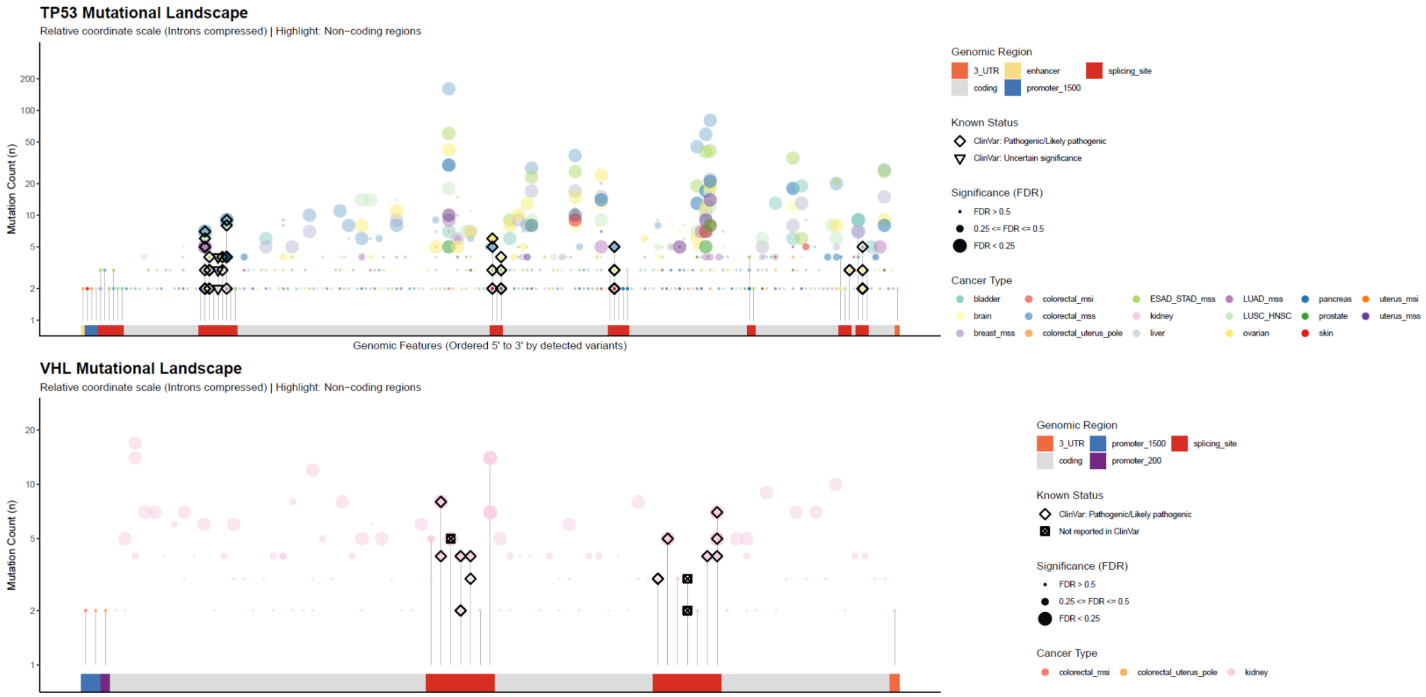


**Supplementary Figure 10. Mutational landscapes of *TP53* and *VHL* splicing and non-coding variants across diverse cancer types.** Lollipop plots illustrating the distribution and observed mutation counts (*n*) of detected variants in *TP53* (top) and *VHL* (bottom). Variants are arranged 5' to 3' across different genomic features with non-coding and splicing regions highlighted. Intronic regions have been compressed using a relative coordinate scale to focus on functional hotspots. Point sizes correspond to False Discovery Rate (FDR) significance categories, and shapes reflect their ClinVar annotation status (e.g., Pathogenic/Likely pathogenic, Uncertain significance, or Not reported). Colors denote the 17 unique molecular and tissue-specific cancer types, highlighting highly recurrent clusters in specific cohorts (such as *VHL* alterations in kidney tumors or multi-cancer *TP53* hotspots).

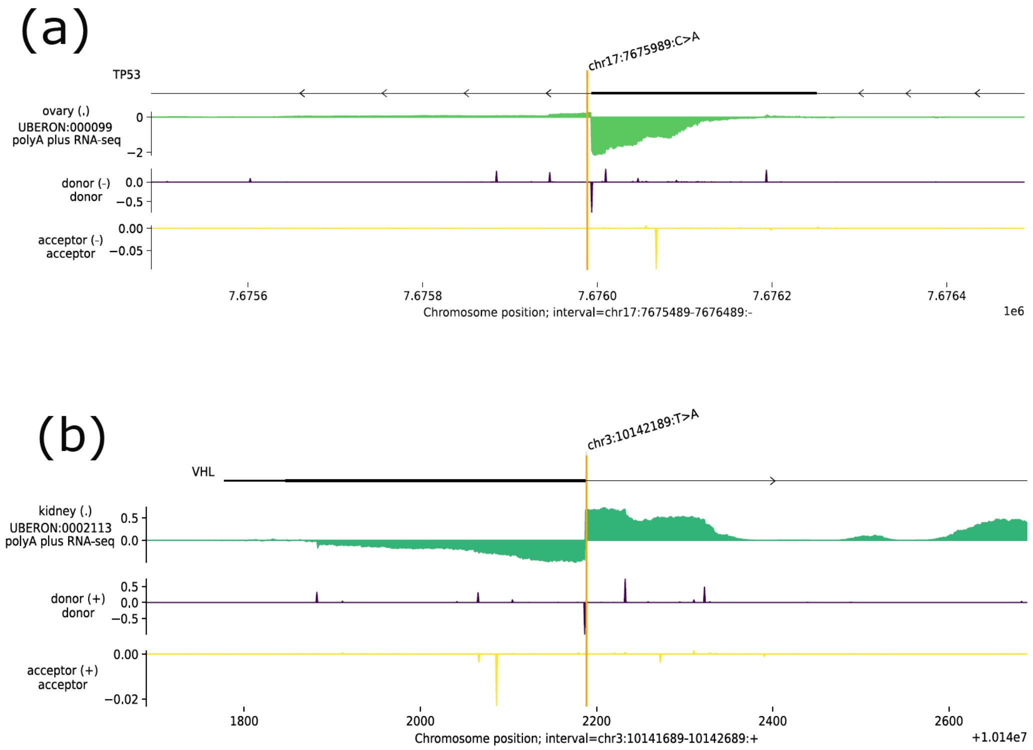


**Supp Figure 11. AlphaGenome genomic tracks and functional predictions for recurrent *TP53* and *VHL* splice region variants.** (a) Recurrent *TP53* splice-region variant. Genomic track visualization surrounding the chr17:7675989:C>A mutation (indicated by the vertical orange line) at the +5 intronic position of TP53 exon 4. Tracks display polyA plus RNA-seq coverage from ovary tissue (green) alongside deep-learning predictions for splice donor (purple) and acceptor (yellow) sites, demonstrating the disruption and abolishment of the canonical donor site. (b) Recurrent VHL canonical splice variant. Genomic track visualization surrounding the chr3:10142189:T>A mutation at the canonical +2 splice site downstream of VHL exon 1. Tracks illustrate baseline kidney RNA-seq coverage (green) coupled with splice donor (purple) and acceptor (yellow) disruption metrics, showing donor site disruption and a predicted extension of the upstream exon.

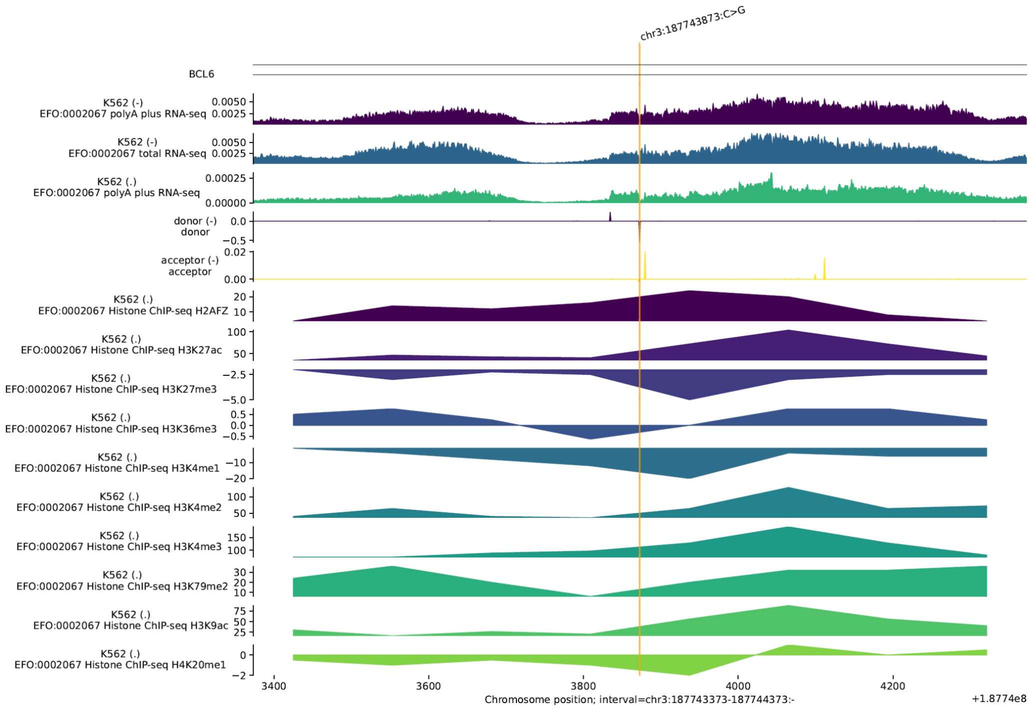


**Supplementary Figure 12. AlphaGenome genomic tracks and functional predictions for the recurrent *BCL6* deep intronic variant.** Computational and epigenetic profile of the chr3:187743873:C>G somatic mutation located +1,537bp from the canonical exon boundary. The upper panels display deep-learning splicing predictions, highlighting a severe disruption of the endogenous splice donor site on the negative strand with a predicted delta score of -0.5. Lower tracks present aligned functional genomic data from the K562 cell line , including polyA plus and total RNA-seq signals , alongside chromatin immunoprecipitation sequencing (ChIP-seq) profiles for key histone modifications (e.g., H3K27ac, H3K4me3, and H3K36me3). These overlapping epigenetic signatures reveal an active chromatin state and robust transcriptional activity at this distal intronic locus, supporting its role as a functional non-canonical splicing driver.

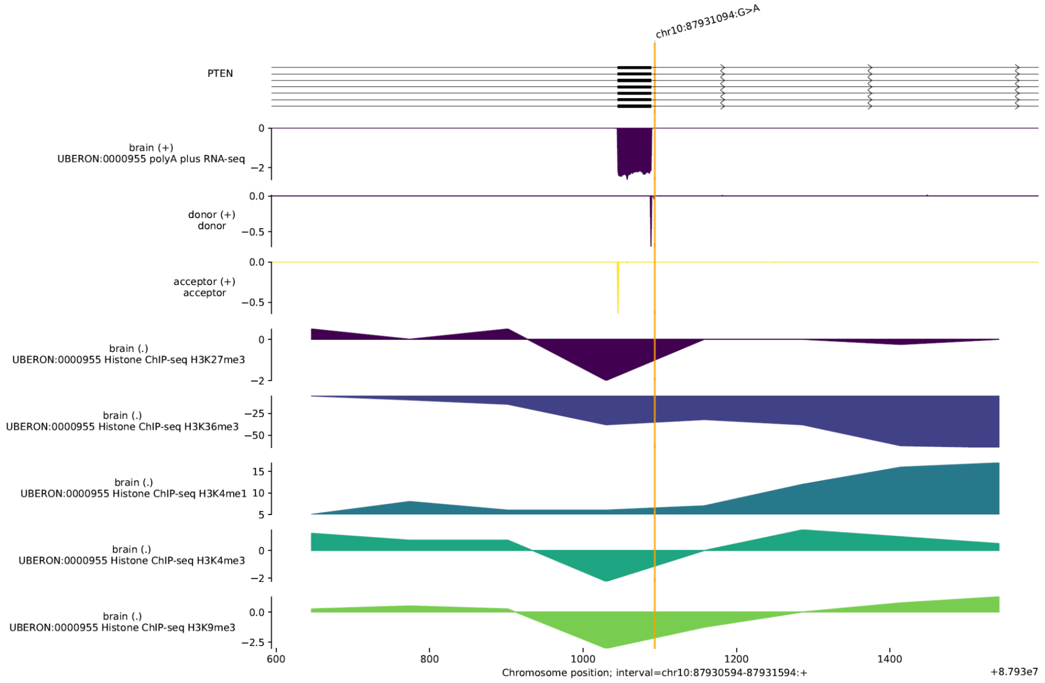


**Supplementary Figure 13. AlphaGenome genomic tracks and functional characterization of a non-canonical *PTEN* splicing hotspot.** Epigenetic and computational splicing architecture flanking the somatic mutation chr10:87931094:G>A, located 5bp away from the canonical exon junction. Upper tracks display splicing prediction models indicating a substantial loss of donor splice site strength on the positive strand, yielding a high-confidence delta score of approximately -0.75, driving a predicted *exon skipping* event. Multi-layered experimental tracks from human brain tissue (UBERON:0000955) demonstrate aligned polyA plus RNA-seq coverage alongside chromatin immunoprecipitation sequencing (ChIP-seq) profiles for active transcription and elongation histone marks (such as robust H3K36me3 signal and low H3K27me3 levels). Despite falling into a marginal statistical bin (0.50 <= FDR < 0.75; $n$ = 3 in brain cancer), the biological significance of this locus is supported by its classification as *Likely Pathogenic* in ClinVar as a germline variant linked to glioma susceptibility phenotypes, strongly supporting its role as a functional somatic driver.

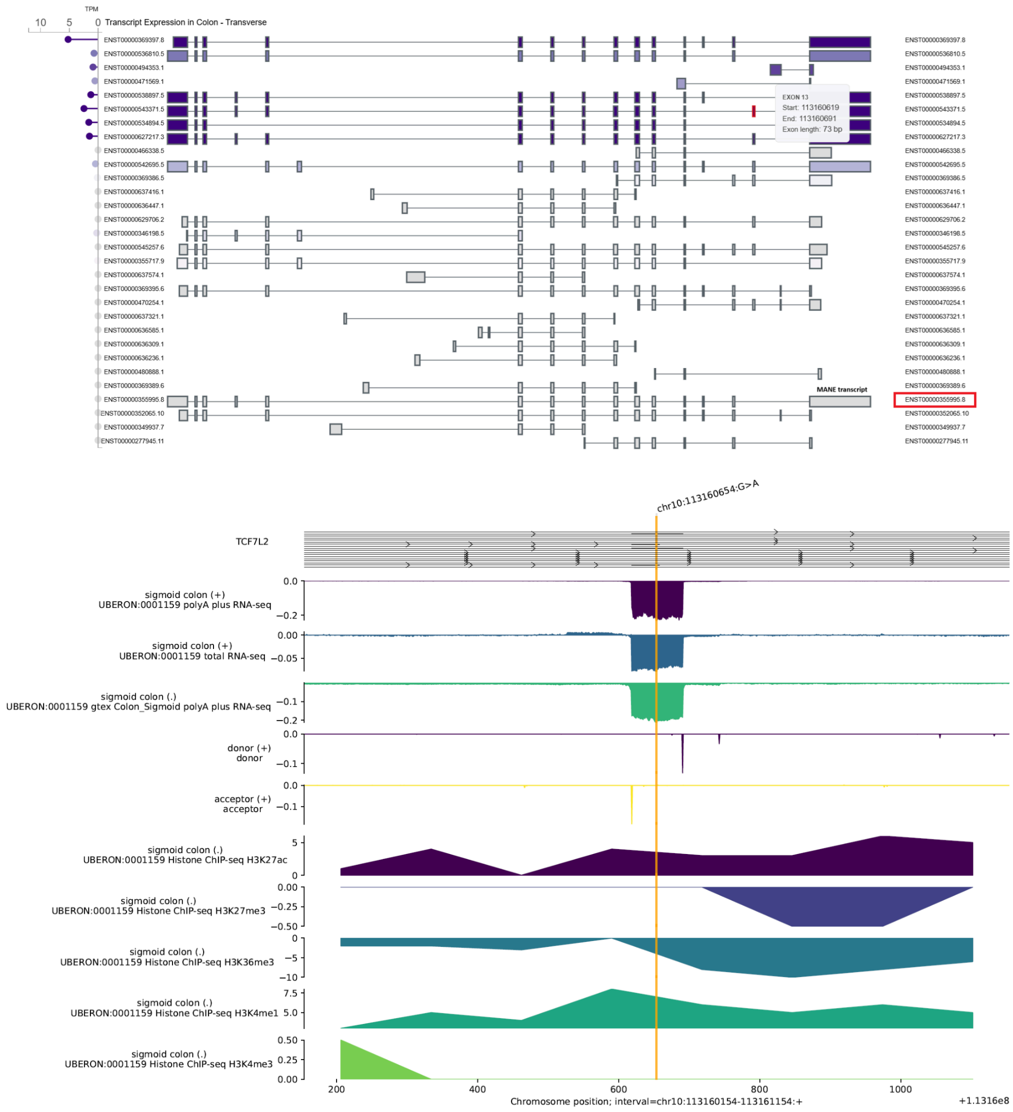


**Supplementary Figure 14. GTEx transcript expression profile of *TCF7L2* in colon tissue and relevance for non-MANE splicing hotspots.** Multi-isoform expression plot from the GTEx database illustrating transcript abundance (Transcripts Per Million, TPM) in transverse colon tissue. The clinically curated MANE transcript (ENST00000355995.8, red box) exhibits negligible or absent expression in this tissue type. Conversely, alternative non-MANE isoforms (e.g., ENST00000369397.8 and ENST00000538897.5, purple tracks) are highly and predominantly expressed. This isoform landscape provides critical biological context for the recurrent colorectal-MSS hotspots at chr10:113160642:C>T (n=7) and chr10:113160654:G>A (n=6). Although classified as "deep intronic" relative to the MANE reference transcript (spanning >650bp away), AlphaGenome (bottom plot) paradoxically predicts *exon skipping* targeted at Exon 13 (73 bp, red marker highlight) of these highly expressed alternative transcripts.

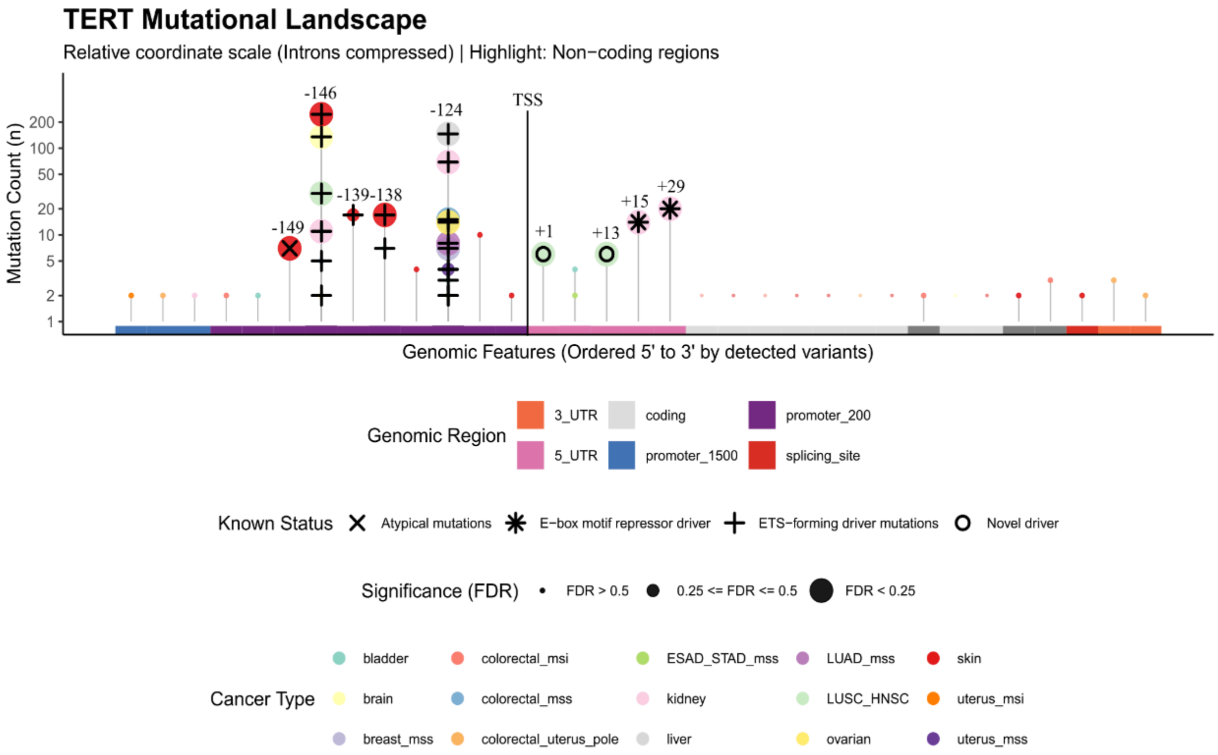


**Supplementary Figure 15. Mutational landscape and positive selection of *TERT* non-coding variants across multiple cancer types.** The lollipop plot displays the frequency and distribution of somatic mutations detected across the *TERT* locus, ordered from 5' to 3' along a relative coordinate scale where intronic regions have been compressed for visualization. The lower genomic track indicates the specific functional annotations, highlighting the core promoter (purple) and the 5' UTR (pink). Each point represents a unique mutated position, where the vertical axis denotes the log10-transformed mutation count (*n*) and individual points are color-coded by their respective cancer type. The size of each circle reflects its statistical significance group based on our mutation recurrence selection (False Discovery Rate, FDR). Canonical core promoter drivers are highlighted as positive controls. Recurrent non-coding mutations within the 5' UTR are explicitly marked, differentiating between previously reported kidney-specific variants and newly identified driver candidates enriched in the LUSC+HNSC (lung squamous cell and head and neck squamous cell carcinoma) cohort; these specific 5' UTR variants disrupt a functional E-box motif, acting as repressor drivers distinct from classic *de novo* ETS-forming promoter mutations.

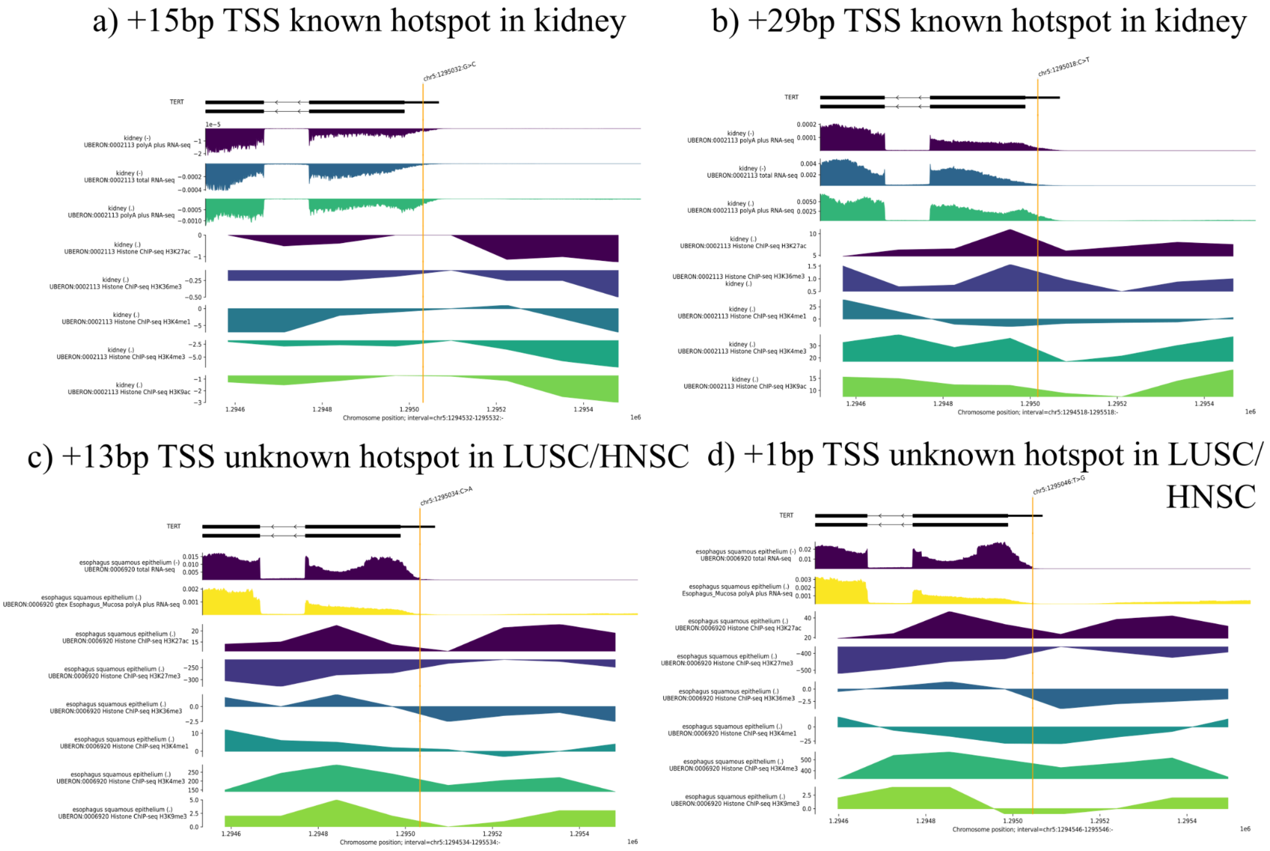


**Supplementary Figure 16. AlphaGenome predictions for the variants in the *TERT* 5' UTR.** AlphaGenome plot displaying the *TERT* 5' UTR mutational landscape, highlighting the previously reported kidney cancer drivers at +15 and +29 bp from the TSS [Mitchell et al., 2018], and the novel LUSC/HNSC hotspots discovered here at +1 and +13 bp distance from the TSS.

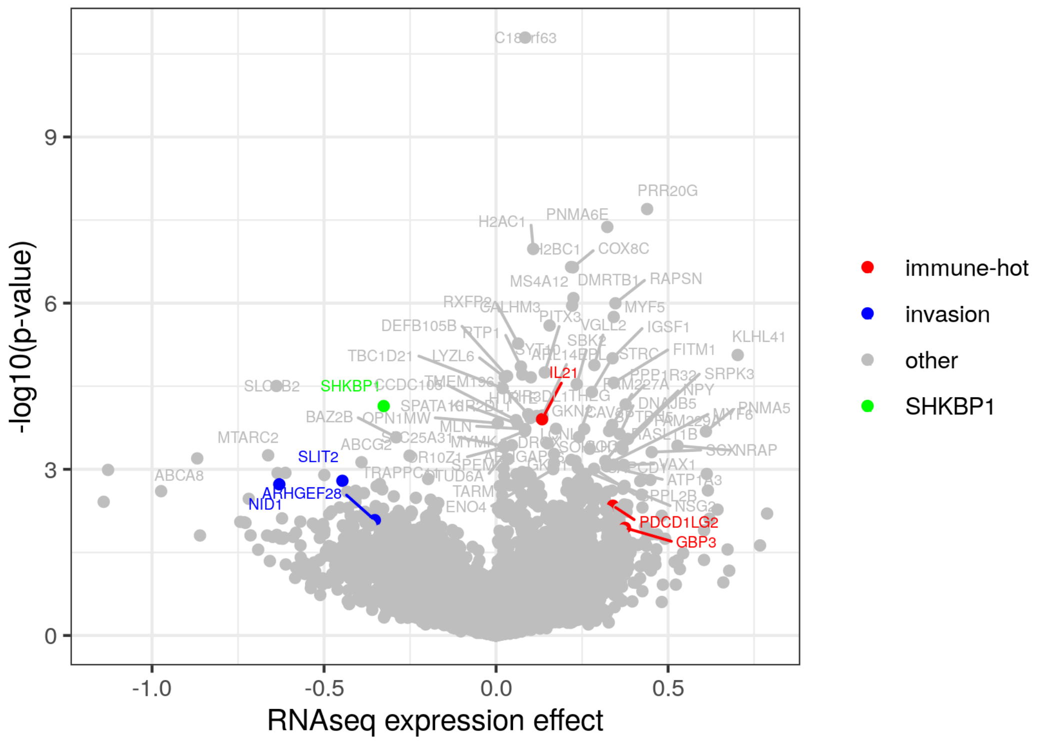


**Supplementary Figure 17. Differential gene expression analysis of *SHKBP1* promoter mutant melanoma.** Volcano plot highlighting significantly differentially expressed genes associated with *SHKBP1* promoter mutations consistently across both the TCGA and HMF melanoma cohorts. Genes driving the "immune-hot" phenotype and heightened invasiveness signatures are labeled. The signature reveals an upregulation of immune-infiltration and invasive markers in mutant samples, providing a molecular context for the counterintuitive increase in EGFR degradation observed in these tumors.

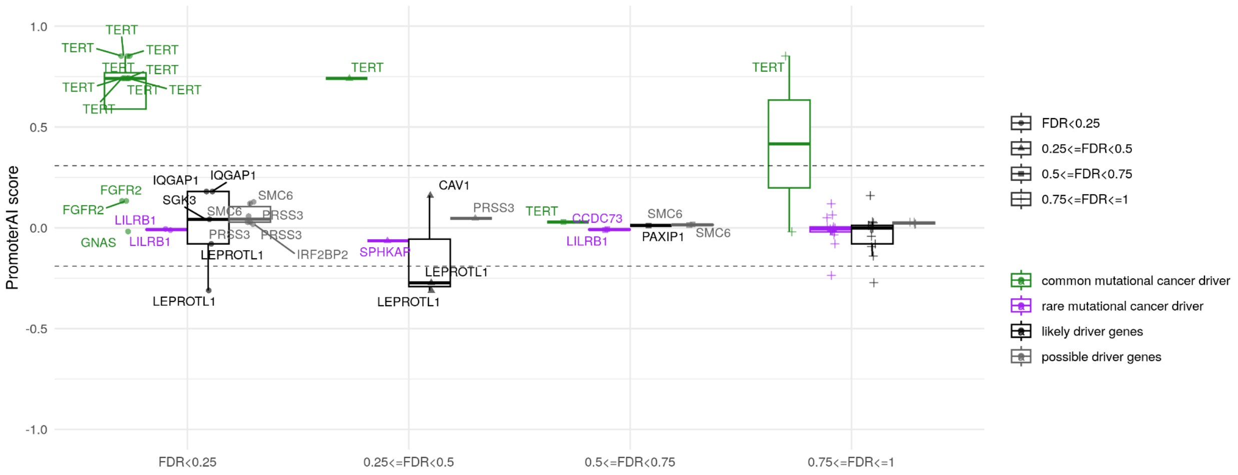


**Supplementary Figure 18. Evaluation of promoter hotspot selection and predicted functional impact across non-skin cancers.** Boxplots illustrate the relationship between mutation recurrence selection (FDR, x-axis) and PromoterAI functional impact scores (y-axis) for TSS-proximal non-coding mutations. Dashed vertical lines represent the operational thresholds for impactful mutations, defined by the 2.5th and 97.5th percentiles of the score distribution in passenger genes.

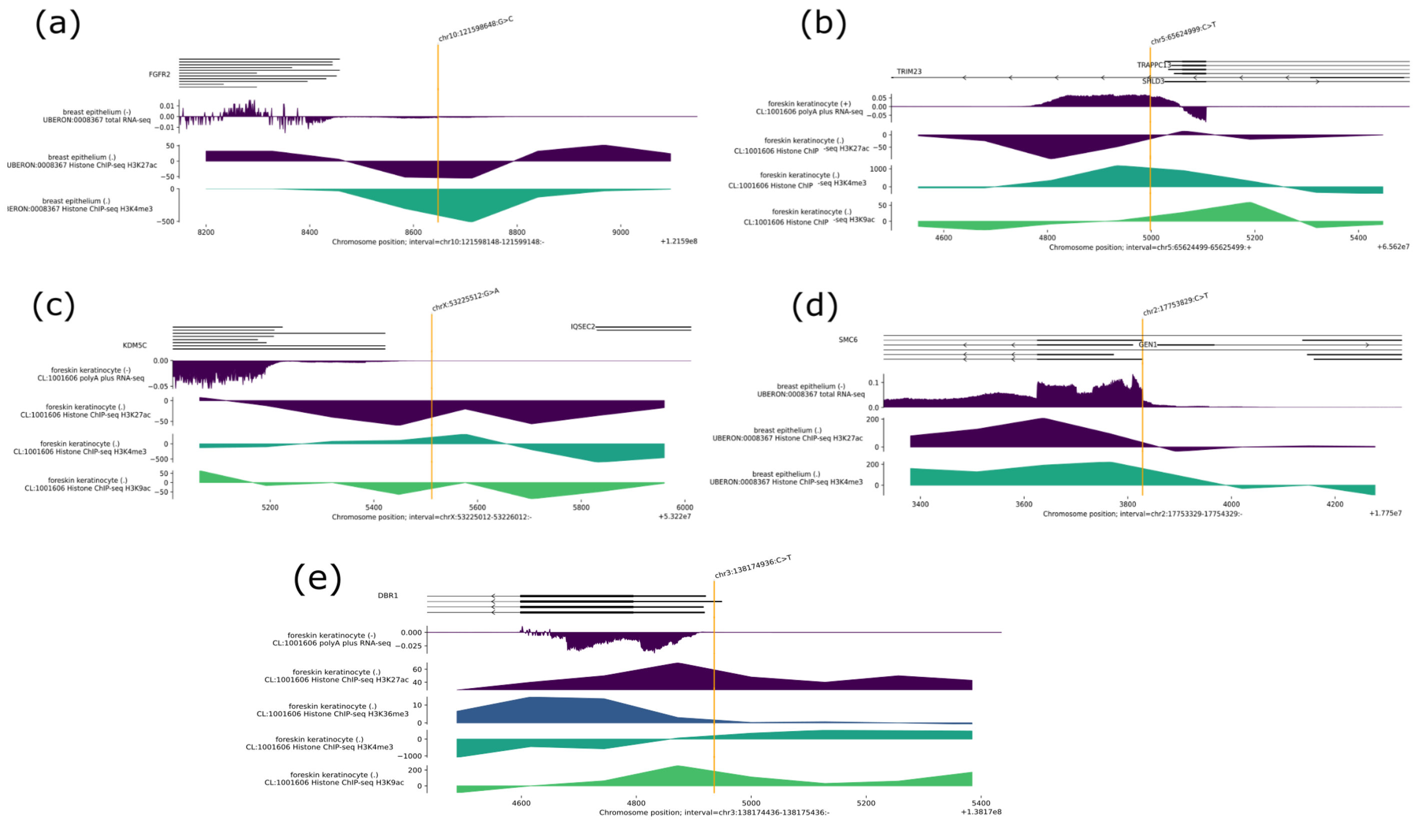


**Supplementary Figure 19. AlphaGenome functional validation of moderate-impact PromoterAI hotspot candidates.** AlphaGenome representations displaying the functional validation of mutated promoter activity across independent models for candidate driver genes approaching the neutral score boundaries: (a) *FGFR2*, (b) *SHLD3*, (c) *KDM5C* (d) *SMC6* and (f) *DBR1*. The plots highlight the variability in promoter activity shifts induced by non-coding somatic variants.

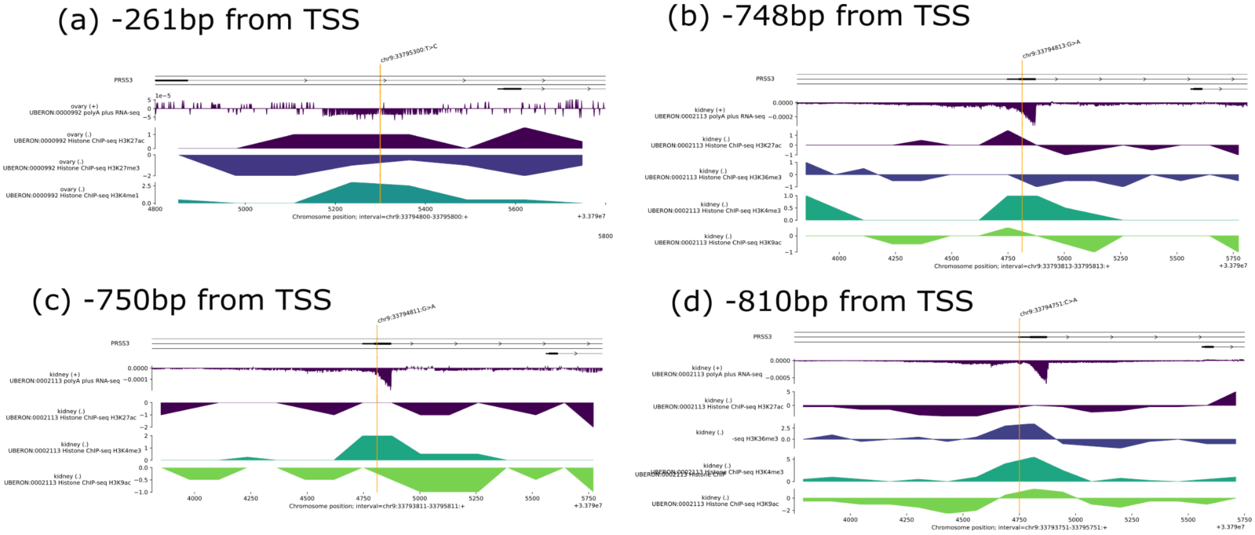


**Supplementary Figure 20. AlphaGenome functional validation of recurrent clustered hotspots in promoter of PRSS3.** AlphaGenome representations displaying the functional validation of mutated promoter activity across recurrent hotspots at (a) -261bp from TSS, (b) -748bp from TSS, (c) -750bp from TSS and (d) -810bp from TSS.

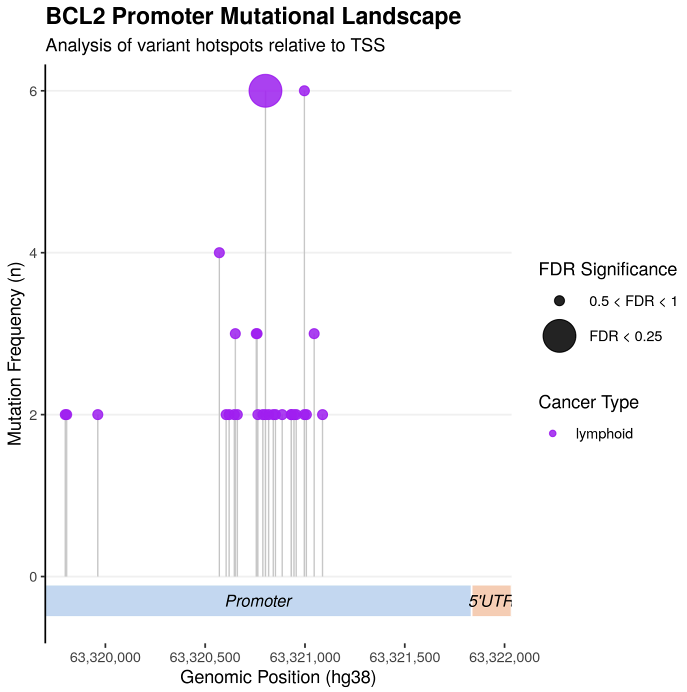


**Supplementary Figure 21. Recurrent hotspots clustered in the promoter of BCL2.** Lolliplot representation displaying the lymphoid-specific mutational landscape across the *BCL2* promoter region, highlighting the 28 recurrent positions (n>=2) and the prominent hotspot at 1,030 bp distal to the TSS (n=6, FDR = 9%).
